# Contrasting patterns of differentiation among three taxa of the rapidly diversifying orchid genus *Ophrys* sect. *Insectifera* (Orchidaceae) where their range overlap

**DOI:** 10.1101/2024.04.23.590674

**Authors:** Pascaline Salvado, Anaïs Gibert, Bertrand Schatz, Lucas Vandenabeele, Roselyne Buscail, David Vilasís, Philippe Feldmann, Joris A. M. Bertrand

**Affiliations:** Laboratoire Génome et Développement des Plantes (LGDP), UMR 5096, Université de Perpignan Via Domitia (UPVD) - Centre National de la Recherche Scientifique (CNRS), Perpignan, France; CEFE, CNRS, Univ Montpellier, EPHE, IRD, Montpellier, France; Centre de Formation et de Recherche sur les Environnements Méditerranéens (CEFREM), Université de Perpignan Via Domitia (UPVD) - Centre National de la Recherche Scientifique (CNRS), Perpignan, France; Grup Orquidològic de Catalunya GOC HCHN; Association Abiome, Olivet, France

**Keywords:** biogeography, genotyping by sequencing, inter-specific gene flow, morphometrics, plants, population genomics, RAD-sequencing, speciation, systematics, taxonomy

## Abstract

In rapidly diversifying groups, taxa defined on the basis of typological criteria can be difficult to support with genetic data. The diversity observed in the insect-mimicking orchid genus *Ophrys* perfectly illustrates this situation; among 400 described species only 9-10 lineages are detectable by genetic markers such as nrITS. The three taxa described in the *Ophrys insectifera* group: *O. insectifera*, *O. subinsectifera* and *O. aymoninii*, can be clearly distinguished by their flowers, which have evolved different phenotypes as a result of adaptation to specific pollinator insect species from three different families. However, genetic differentiation between these three taxa has never been really supported by population genetic data and their taxonomic status is still debated. Using population genomic approaches, we found a clustering consistent with the existence of three genetic entities where the geographic distributions of the three taxa overlap. Two of these clusters correspond to France populations of the widespread *O. insectifera* and the micro-endemic *O. aymoninii*. However, the last cluster grouped together all the Iberian individuals, suggesting that individuals phenotypically identified as either *O. insectifera* or *O. subinsectifera* are genetically weakly differentiated there. Populations of the two pairs of taxa thus may have experienced different patterns of inter-specific gene flow.

## Introduction

Adaptive radiations can lead to rapid accumulation of species through adaptation to various selection pressures. In such cases, reconstructing the evolutionary history based solely on genetic data can be challenging. The advent of high-throughput sequencing technologies has enabled to work at the whole-genome scale, thus providing significant gain in the resolution of genetic data, even in non-model organisms (e.g., Léveillé-Bourret *et al*. 2020). For species with large and/or complex genomes however, genome complexity reduction protocols remain the most efficient way to simultaneously obtain genomic data on a sufficient number of individuals and populations in order to investigate how evolutionary lineages diverge from each other.

*Ophrys* orchids are renowned for their exceptional ability to attract male insect pollinators through their insect-mimicking flowers. The labellum, one of the petals, is especially effective in luring the insects to the point where they attempt to copulate with it, ultimately ensuring the plant’s reproduction. The degree of specification required for the emergence and the maintenance of this plant-insect interaction is likely to be responsible for the high rate of diversification observed in *Ophrys* orchids (see Baguette *et al*. (2020) for a recent review). The flower phenotypes of *Ophrys* orchids are subject to strong selection pressures exerted by male pollinator insects, which is one of the main driver of this extreme adaptive radiation. The morphological diversity observable among *Ophrys* species has been well-documented and an increasing number of studies have reported subtle variations in flower odor compounds associated with pollinator species identity (e.g. Schlüter *et al*. 2011; Joffard, Buatois and Schatz 2016; Gervasi *et al*. 2017). However, genetic studies have often failed to characterise patterns and propose relevant scenarios to explain evolutionary divergence between closely related *Ophrys* species.

The inability of traditional genetic markers to delineate *Ophrys* lineages can be explained by divergence times that may be too recent to display allele sorting through genetic drift (*i.e.* Incomplete Lineage and Sorting) and/or reticulate evolution that may blur any emerging phylogenetic signal. In spite of the promising results from a pioneering study using a genotyping-by-sequencing (GBS) protocol to investigate patterns of differentiation between closely related species of the *Ophrys sphegodes* group (Sedeek *et al*. 2014), population genomic approaches have not been used in *Ophrys* since. Bateman *et al*. (2018) have successfully used RADseq-like protocol data to clarify the phylogenetic relationships between the main *Ophrys* lineages, see also Piñeiro Fernández *et al*. (2019) who used RNAseq data and Bertrand *et al*. (2021) who employed whole plastomes. However, none of these studies addressed patterns of species differentiation at lower taxonomic scale (but see Bateman *et al*. 2021). Thus, the actual taxonomic status of the so-called ‘microspecies’ of *Ophrys* is still regularly contested (e.g. Bateman and Rudall 2023) although the suitability of population genomic approaches to investigate evolutionary divergence between *Ophrys* lineages has rarely been properly tested. It was recently confirmed that a population-based sampling design coupled with an appropriate genome subsampling protocol (*i.e.* a genotyping-by-sequencing protocol called nGBS, which stands for normalized Genotyping-By-Sequencing) allowed to accurately characterize patterns of diversity and differentiation among little divergent populations and taxa of *Ophrys aveyronensis* (Gibert *et al*. 2023).

In this study, we used a protocol similar to the one of Gibert *et al*. 2023 to test whether this approach also provides sufficient resolution to distinguish taxa within another *Ophrys* clade whose systematics and taxonomy are problematic: the *Ophrys insectifera* group. The latter consists of three described taxa: the Euro-Mediterranean widespread *Ophrys insectifera* L. and two endemic taxa: *Ophrys subinsectifera* (Hermosilla and Sabando 1996), whose repartition spans the Southern flanks of the Pyrenees and *Ophrys aymoninii* (Brestr.) Buttler, which is (micro-)endemic to the Grand Causses region in France. These three taxa currently recognized as species or subspecies (*Ophrys insectifera* subsp. *subinsectifera* (Hermosilla and Sabando 1996) (O.Bolòs & Vigo) and *Ophrys insectifera* subsp. *aymoninii* Breistr.) are known to be pollinated by insect species belonging to three different families, have been shown to display differences in odor bouquet and use to be easily distinguishable in the field based on their phenotypes. So far however, genetic studies have failed to delineate these taxa based on genetic data (Triponez *et al*. 2013; Gervasi *et al*. 2017). Previous phylogenetic studies only included one or two individuals per taxon often with only two of the three taxa of the group, which did not allow to properly test their reciprocal monophyly (Soliva and Widmer 2003; Devey *et al*. 2008; Breitkopf *et al*. 2015; Bateman *et al*. 2018). With the future aim of investigating the ecological and evolutionary causes of species boundaries in this system, we tested whether population genomic data could allow to characterize species, especially where *O. insectifera* co-occur with one of the two endemics. Specifically, we tested whether population genomics allow to distinguish the three taxa despite incomplete lineage sorting and possible interspecific gene flow. If so, we planned to assess for the first time levels of genetic diversity within taxa and genetic differentiation between taxa based on genomic data.

## Material & Methods

### Population sampling

We sampled a total of 239 individuals at 12 localities in May 2022 and 2023, where the widespread *O. insectifera* locally co-occurs with one of the endemic taxa: *O. subinsectifera* in Catalonia, Spain at 6 sites (1-Les Guilleries, 2-Puig Grifó, 3-Tarradell, 4-Castanyola, 5-L’Estanyol and 6-Seva) and *O. aymoninii* in the Grands Causses Region, France at 4 sites (8-Le Buffre, 9-Col de Montmirat, 10-Nivoliers and 11-Fretma) as well as at two additional localities in between these two regions where only *O. insectifera* is present (7-Versols-et-Lapeyre and 12-Rodome), as represented in Fig.1 (see also Table 1 and Supplementary Table S1). At each locality, we collected ca. 1-2 cm^2^ of leaf tissues that were stored in 90% ethanol at 4°C until DNA extraction. *O. subinsectifera* and *O. insectifera* are not protected in the study area but *O. aymoninii* has a Near Threatened status (NT) on the French National Red List. We asked for and obtained special permits from the “Parc National des Cévennes” to collect plants and drive within the national park (2022-0115, 2022-0117, 2023-0093 and 2023-0094). The research project also complies with the Nagoya protocol (TREL2206915S/562), which provides a framework for the fair and equitable sharing of benefits arising from the utilization of genetic resources. We photographed each individual and obtained phenotypic values at a suite of ten phenotypic traits, as described in detail in Gibert *et al*. (2022). We recorded five “whole plant” traits: plant size, stem diameter, number of reproductive structures (flowers and buds), distance from the ground to the first flower, and distance between the first and the second flower. We also measured five floral traits: length and width of the labellum and the median sepal as well as the length of the right petal. We calculated the ratio between the length and width of the labellum to get an overall idea of its proportions. All the individuals were categorized as *Ophrys insectifera*, *O. aymoninii*, *O. subinsectifera* or intermediates (*O. aymoninii*/*O. insectifera* or *O. subinsectifera*/*O. insectifera*) based on established visual criteria (*i.e.* following Delforge 2021). We performed LMM (Linear Mixed Models) on each traits to test for difference between the five categories. The fixed effect included the categories and the random effet included the study site. We presented 95% confidence intervals, and used their lack of overlap with zero or lack of overlap between categories to define what effets were statistically significant, in addition to *p*-values. This is a conservative approach that provides information on the statistical effect (range, direction, strength and reliability) that is not provided by *p*-values (Ho *et al*. 2019). Models were implemented in R via the ‘lme4’ package (Bates, Martin and Walker 2016). Assumptions of models (e.g. heteroscedasticity, over and under dispersion, significance of outliers) were checked both visually and by using the DHARMa R-package (Hartig 2018).

**Figure 1.**
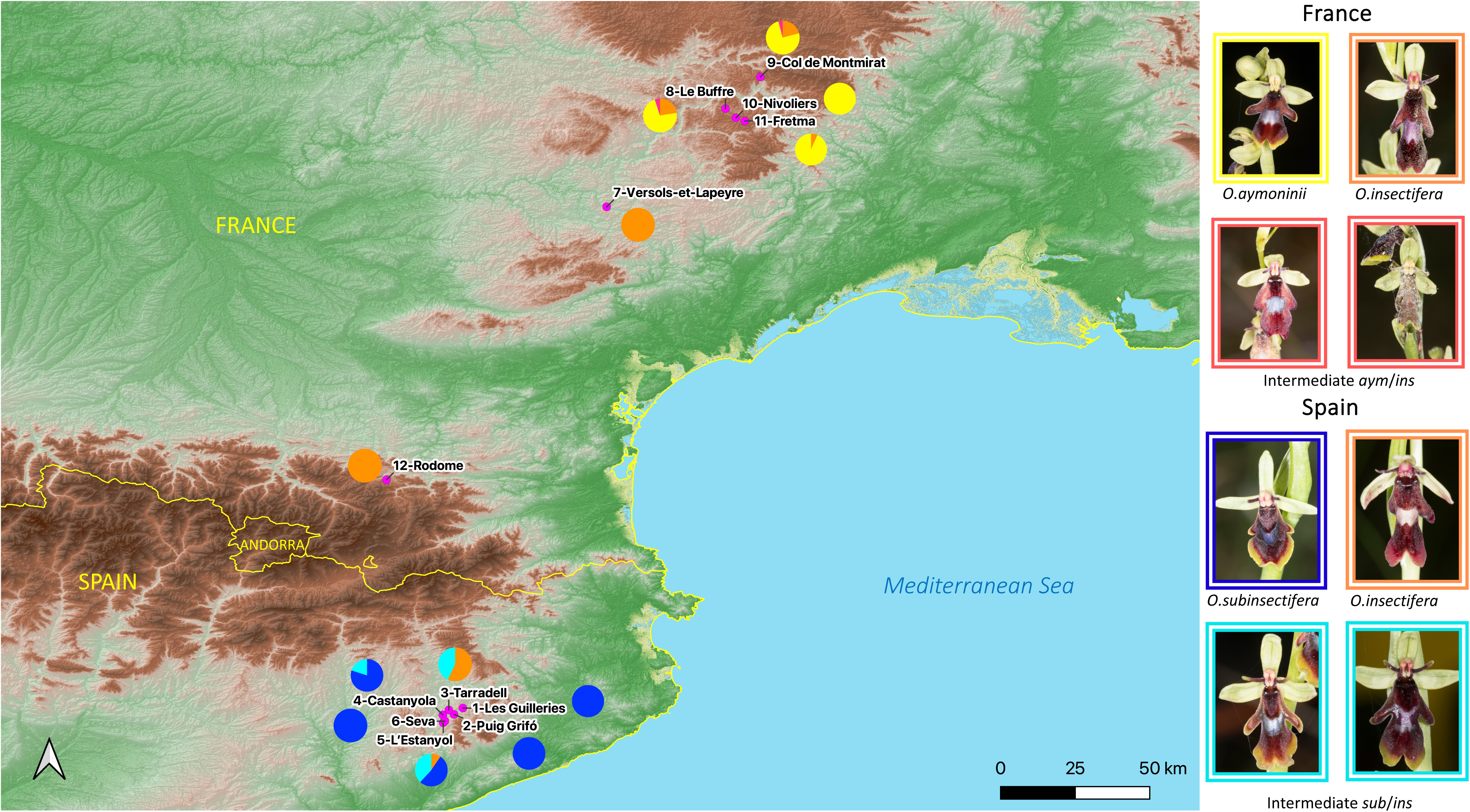
Map displaying the 12 study sites: 4 in France for *O. aymoninii*, *O. insectifera* and their intermediates (8-Le Buffre, 9-Col de Montmirat, 10-Nivoliers and 11-Fretma), 2 “control” sites in France with only *O. insectifera* (7-Versols and 12-Rodome) and 6 sites in Spain for *O. subinsectifera*, *O. insectifera* and their intermediates (1-Les Guilleries, 2-Puig Grifó, 3-Tarradell, 4-Castanyola, 5-L’Estanyol and 6-Seva). Pie charts represent the species composition (on the basis of identification on the basis of phenotypes) of each site, yellow corresponding to *O. aymoninii*, orange to *O. insectifera*, dark blue to *O. subinsectifera*, red for the intermediates *aym*/*ins* and light blue for the intermediates *sub*/*ins*.

**Table 1.** Geographic coordinates (in °) and elevation (in m above sea level) of the sampling localities, sample size (*N*), observed (*H*_O_) and expected (*H*_E_) levels of heterozygosity, deviation from panmixia (*G*_IS_), total number of alleles (*A*) and number of private alleles (*A*p) per locality.

| Locality | Latitude | Longitude | Elevation | $N$ | $H_O$ | $H_E$ | $G_{IS}$ | $A$ | $A_p$ |
| --- | --- | --- | --- | --- | --- | --- | --- | --- | --- |
| 1-Les Guilleries | 41.883 | 2.356 | 602 | 30 | 0.228 | 0.254 | 0.103 | 11158 | 2 |
| 2-Puig Grifó | 41.857 | 2.320 | 786 | 20 | 0.226 | 0.249 | 0.096 | 11034 | 0 |
| 3-Tarradell | 41.876 | 2.300 | 681 | 7 | 0.229 | 0.224 | -0.023 | 10750 | 0 |
| 4-Castanyola | 41.855 | 2.276 | 604 | 15 | 0.233 | 0.247 | 0.058 | 10980 | 0 |
| **5-Seva** | 41.824 | 2.277 | 742 | 21 | 0.232 | 0.245 | 0.052 | 10756 | 0 |
| **6-L'Estanyol** | 41.831 | 2.284 | 715 | 24 | 0.236 | 0.244 | 0.036 | 10890 | 1 |
| **7-Versols-et-Lapeyre** | 43.897 | 2.934 | 412 | 17 | 0.232 | 0.267 | 0.131 | 11066 | 0 |
| **8-Le Buffre** | 44.292 | 3.411 | 922 | 40 | 0.233 | 0.272 | 0.143 | 11182 | 0 |
| **9-Col de Montmirat** | 44.419 | 3.551 | 1068 | 24 | 0.226 | 0.253 | 0.105 | 10846 | 0 |
| **10-Nivoliers** | 44.255 | 3.453 | 1017 | 14 | 0.225 | 0.266 | 0.155 | 10679 | 0 |
| **11-Fretma** | 44.242 | 3.389 | 1117 | 15 | 0.235 | 0.232 | -0.013 | 10868 | 4 |
| **12-Rodome** | 42.800 | 2.049 | 959 | 12 | 0.241 | 0.267 | 0.095 | 10825 | 8 |
| **Mean** |  |  |  |  | **0.231** | **0.252** | **0.078** |  |  |

### Molecular procedures

Genomic DNA extraction and genotyping were subcontracted to LGC Genomics GmbH (Berlin, Germany) following a protocol called nGBS for ‘normalised Genotyping By Sequencing’. This double digest Restriction site-Associated DNA seq or RAD-seq-like protocol relies on two restriction enzymes (PstI and ApeKI in our case) to reduce *Ophrys* genome complexity and it includes a normalization step that aims at avoiding the overrepresentation of repetitive regions. The resulting 239 individually barcoded libraries were sequenced in paired-end mode (2 x 150 bp) on an Illumina NovaSeq 6000 with an expectation of a minimum number of 1.5 million read pairs per sample. We used Stacks v.2.62 (Catchen *et al*. 2011, 2013) to build loci from Illumina reads. After demultiplexing and cleaning the samples with *process_radtags*, we aligned them to the reference genome of *Ophrys sphegodes*, which is the most closely related reference genome available (Russo *et al*. 2023) using BWA-mem v.0.7.17 (Li and Durbin 2009). Then, *ref_map.pl* pipeline was executed, for building RAD-loci and SNP calling. Finally, we ran the *populations* script included in Stacks, from which only loci that were genotyped in at least 80% of the individuals per population (-*r 80*) and 10 populations (out of 12) were kept (-*p 10*), while loci exhibiting estimates of observed heterozygosity greater than 70% (-- *max-obs-het 0.7*) were filtered out to reduce the risk of including remaining paralogs. We also discarded sites whose minor allele frequency was lower than 5% (--*min-maf 0.05*). The genotype matrix of the remaining 11,089 SNPs was then exported in several formats such as Structure, VCF, genepop, radpainter.

### Population genomics analyses

To get an overview of the overall genetic diversity and differentiation among individuals and populations, we first performed a Principal Component Analysis (PCA) based on a matrix of 239 individuals (as rows) and 11,089 SNPs (as columns) coded as a *genlight* object with the R-package ‘adegenet’ (Jombart and Ahmed 2011). We then used Genodive v.3.04 (Meirmans 2020) to compute expected and observed heterozygosity (*H*_E_ and *H*_O_, respectively). For all the analyses realized in Genodive, we kept only the most abundant species at each site. The number of alleles (*A*) and private alleles (*A*_P_) were retrieved from the *populations* outputs given by Stacks. The deviation from panmixia was evaluated by computing *G*_IS_ and the statistical significance of the obtained values was estimated based on 10,000 permutations, in Genodive. Overall and pairwise genetic differentiation was assessed based on *G*_ST_, (Nei 1987) as implemented in Genodive in a similar manner. Genodive was also used to carry out a hierarchical Analysis of Molecular Variance (AMOVA) to assess proportions of genetic variance (i) among series (here, Series A between *O. aymoninii* and *O. insectifera* and Series B between *O. subinsectifera* and *O. insectifera*) (*F*_CT_) and (ii) among populations within Series (*F*_SC_). The statistical significance of the obtained values was estimated based on 10,000 permutations. To further investigate population structure and characterize putative migration/admixture event, we used ADMIXTURE (Alexander, Novembre and Lange 2009). This method computes ‘*ancestry coefficients’* that represent the proportions of an individual genome that originate from multiple ancestral gene pools. The number of genetic clusters was varied from *K* = 1 to 12, analyses were run with 10 replicates and a cross-validation procedure was performed to determine the optimal value of *K*. To assess genetic co-ancestry among individuals within and across the different sites, we used haplotype-based population inference approach implemented in *fineRADstructure* v.0.3.1 (Malinsky *et al*. 2018) using the same filtering criteria described for the SNP dataset. Analyses were performed using default settings and results were visualized with dedicated R-scripts provided with the program.

## Results

### Species distribution

In both France and Spain, we sampled sites with only the endemics (*O. aymoninii* or *O. subinsectifera*), others with only the widespread *O. insectifera*, and also sites with both one of the endemics and *O. insectifera* in close proximity (Fig. 1). In Spain, sites 3-Tarradell, 4-Castanyola and 5-L’Estanyol showed the presence of *O. insectifera*, *O. subinsectifera*, and a non-negligible number of individuals with intermediate phenotypes (15 out of 117, 13%), which could not be clearly assigned to one species or the other. Conversely, sites 1-Les Guilleries, 2-Puig Grifó and 6-Seva exclusively presented individuals with phenotypes corresponding to *O. subinsectifera*. In France, our survey revealed three sites (8-Le Buffre, 9-Col de Montmirat and 11-Fretma) where *O. aymoninii* and *O. insectifera* co-occurred. However, we found only a few individuals with intermediate phenotypes (3 out of 123, 2.4 %) at sites 8-Le Buffre and 9-Col de Montmirat (we did not observe *O. insectifera* at site 10-Nivoliers). At sites 7-Versols-et-Lapeyre and 12-Rodome, we observed only *O. insectifera*, which makes sense as these two sites are outside the known range of both endemic species. Therefore, these sites were considered as unadmixed with either one of the endemics and served as reference sites for *O. insectifera* in our study.

### Patterns of phenotypic variation

For morphometric analyses, we considered n= 50 *O. insectifera* individuals from sites 3-Tarradel, 5-L’Estanyol, 7-Versols-et-Lapeyre, 8-Le Buffre, 9-Col de Montmirat, 11-Fretma and 12-Rodome, n=95 *O. subinsectifera* individuals from sites 1-Les Guilleries, 2-Puig Grifó, 4-Castanyola, 5-L’Estanyol and 6-Seva, n=75 *O. aymoninii* individuals from sites 8-Le Buffre, 9-Col de Montmirat, 10-Nivoliers and 11-Fretma, n=14 intermediate individuals between *O. insectifera* and *O. subinsectifera* from sites 3-Tarradell, 4-Castanyola and 5-L’Estanyol and n=3 intermediate individuals between *O. insectifera* and *O. aymoninii* from sites 8-Le Buffre and 9-Col de Montmirat.

Our results confirm that the flowers of the three taxa show differences in morphometrics, with the endemics being the smallest overall. The labellum length of the two endemic species (*O. aymoninii* and *O. subinsectifera*) is significantly shorter (9.4 ± 1.2 mm and 10.6 ± 1.3 mm respectively, mean ± SD, *p*<0.01, Supporting Information Fig. S2 and Table S2) compared to *O. insectifera* (13 ± 1.7 mm, Supporting Information Fig. S2 and Table S2). The labellum ratio differs between the three species; *O. aymoninii* showed a ratio 42% smaller than *O. insectifera* (respectively ratio = 1.4 vs 2.4) and *O. subinsectifera* showed a ratio 15% smaller than *O. insectifera* (2, Supporting Information Fig. S2 and Table S2). Petal length is significantly shorter for *O. subinsectifera* (2.7 ± 0.9 mm, *p*<0.01) and *O. aymoninii* has the shortest sepal length (5 ± 0.8 mm, *p*<0.01). *O. subinsectifera* has a narrower sepal width compared to *O. insectifera* (2.3 ± 0.6 mm, *p*<0.01 Supporting Information Fig. S2 and Table S2). Looking at whole plant traits, we observe that both endemics tend to have smaller plant size compared to *O. insectifera* (19.4 ± 6.4 cm, *p* = 0.01 for *O. subinsectifera* and 19.7 ± 6.3 cm, *p*<0.01 for *O. aymoninii* vs 25.4 ± 5.9 cm for *O. insectifera* Supporting Information Fig. S2 and Table S2). In addition, endemics have more compact inflorescences, whereas those of *O. insectifera* appear looser, with greater distances between the ground and the first flower (16.8 ± 4.9 cm, *p*<0.01, Supporting Information Fig. S2 and Table S2) and between the first and second flower (2.9 ± 0.8 cm, *NS,* Supporting Information Fig. S2 and Table S2). Intermediate individuals, termed “*sub*/*ins*” and “*aym*/*ins*”, show intermediate values for labellum length, labellum ratio, petal length and sepal width (Supporting Information Fig. S2 and Table S2). In some cases, they also tend to have larger plant sizes and stem diameters compared to the three other categories, as well as greater distances between the first and second flower (*p*<0.05, Supporting Information Fig. S2 and Table S2).

### Population genomic dataset

We obtained a total of 1 051 266 532 of read pairs across all the individuals (496 296 - 7 535 054 of raw reads per ind., mean 4 380 277, SD = 1 044 261) and 357 155 reads (< 1%) were removed because of low quality. We genotyped a total of 9 766 159 loci (composed of 2 065 743 449 sites) including 794 414 variant sites (SNPs) with *Stacks*. Average read depth ranged from 7X to 102.7X (mean 63.6X, SD = 16.1X) based on the combination of parameters we used. After filtering the data with populations, we finally kept 13 772 loci from which we retained 11 089 variant sites (SNPs).

### Patterns of genetic diversity and differentiation

All sites were found to have similar levels of genetic diversity (*A* and *A*_R_). The inbreeding index (*G*_IS_) showed that the deviation from panmixia was higher in the French sites, with the exception of site 11-Fretma, compared to Spain (Table 1). The negative *G*_IS_ values for site 3-Tarradell are considered unreliable as they were calculated based on the basis of only four individuals of *O. insectifera*. The AMOVA highlights a higher genetic differentiation between *O. insectifera* from the control sites (7-Versols-et-Lapeyre and 12-Rodome) and *O. aymoninii* from sites 8-Le Buffre, 9-Col de Montmirat, 10-Nivoliers and 11-Fretma with a *F*_CT_ of 0.072 compared to *O. insectifera* from the control sites and *O. subinsectifera* from sites 1-Les Guilleries, 2-Puig Grifó and 6-Seva which displays a *F*_CT_ of 0.038 (Table S3).

Principal Component Analysis (PCA), with PC1 and PC2 explaining 62.49% and 20.11% of the total genetic variance respectively (Fig. 3A), shows that all individuals morphologically assigned to *O. aymoninii* cluster together (although individuals from site 11-Fretma were slightly separated from those from the other sites 8-Le Buffre, 9-Col de Montmirat and 10-Nivoliers). In addition, all individuals assigned to *O. insectifera* from France (at sites 8-Le Buffre, 9-Col de Montmirat, 10-Nivoliers, 11-Fretma) cluster together with the reference sites of 7-Versols-et-Lapeyre and 12-Rodome. However, the individuals morphologically assigned to *O. insectifera* from Spain do not cluster with *O. insectifera* from France. Instead, they were found to cluster with other individuals from Spanish sites. Indeed, in Spain the pattern is not as clear as in France since we can observe two subclusters. The first one groups the sites 1-Les Guilleries, 2-Puig Grifó, 6-Seva, four individuals from site 5-L’Estanyol (*i.e*., 22-Div-025, 22-Div-026, 22-Div-027, 22-Div-044) and 10 individuals from site 4-Castanyola (*i.e*., 22-Div-068, 22-Div-070, 22-Div-071, 22-Div-073, 22-Div-075, 22-Div-076, 22-Div-077, 22-Div-078, 22-Div-079, 22-Div-080) corresponding to *O. subinsectifera*. The second subcluster groups the remaining individuals from sites 4-Castanyola and 5-L’Estanyol, and all the individuals from site 3-Tarradell, corresponding to intermediate individuals between *O.insectifera* and *O. subinsectifera*. Removing sites containing both *O. insectifera* and *O. subinsectifera* and/or intermediate individuals (*i.e*., 3-Tarradell, 4-Castanyola and 5-L’Estanyol) and leaving sites containing only individuals with *O. subinsectifera* phenotypes, results in an even clearer clustering of the species into three entities (with PC1 and PC2 explaining 62.28% and 20.56% of the total genetic variance respectively), this time corresponding to three species *O. subinsectifera*, *O. insectifera* and *O. aymoninii* (Fig. 3B).

**Figure 2.**
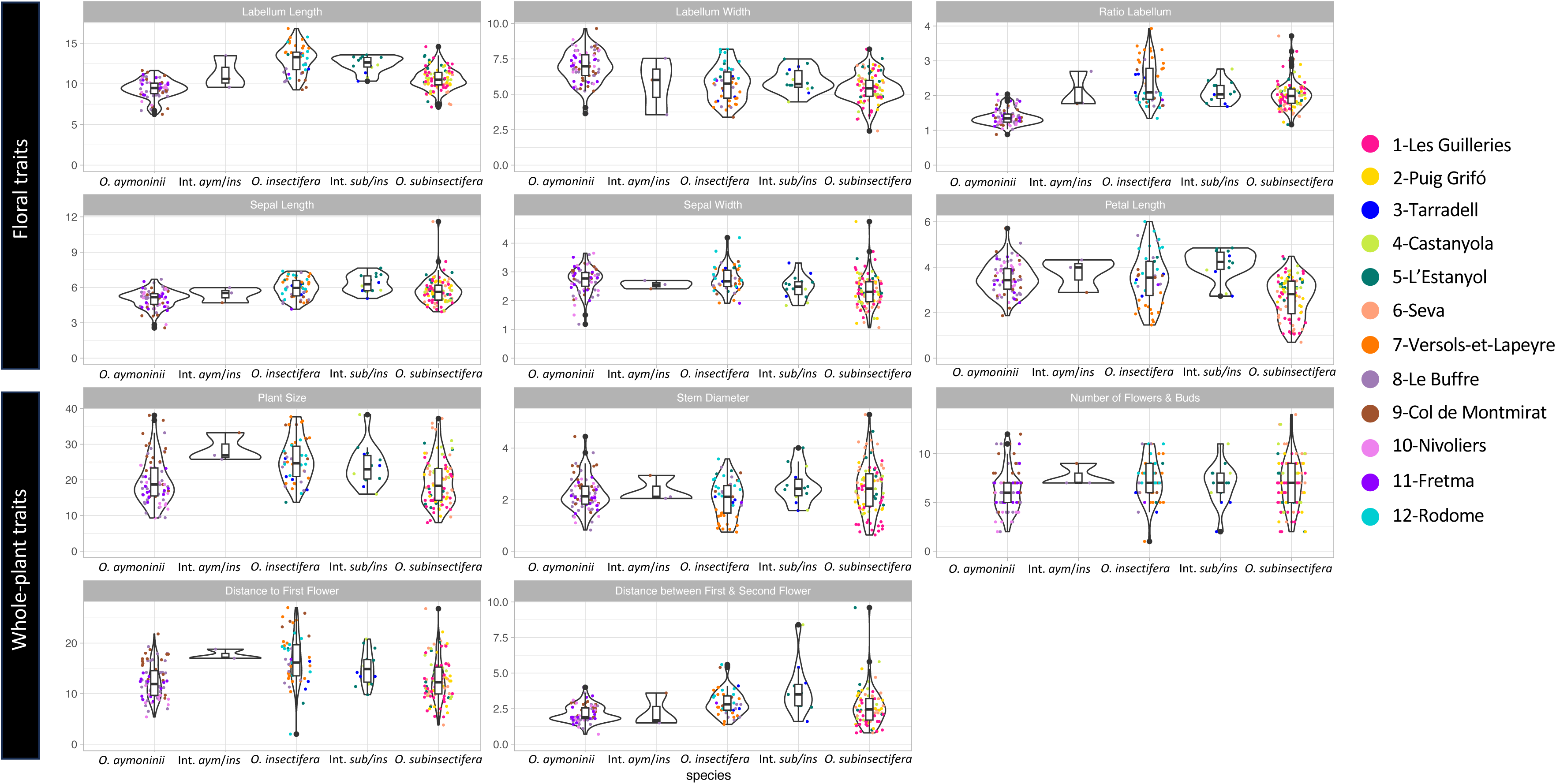
Distribution (violin plots and boxplots) of 6 floral and 5 whole-plant traits for each species: *O. aymoninii*, *O. insectifera*, *O. subinsectifera* and individuals with intermediate phenotypes: *aym*/*ins* and *sub*/*ins*. The different sites are displayed in different colors. All the values are in mm except for the Plant Size, Distance to First Flower and Distance between First and Second Flower which are in cm.

**Figure 3.**
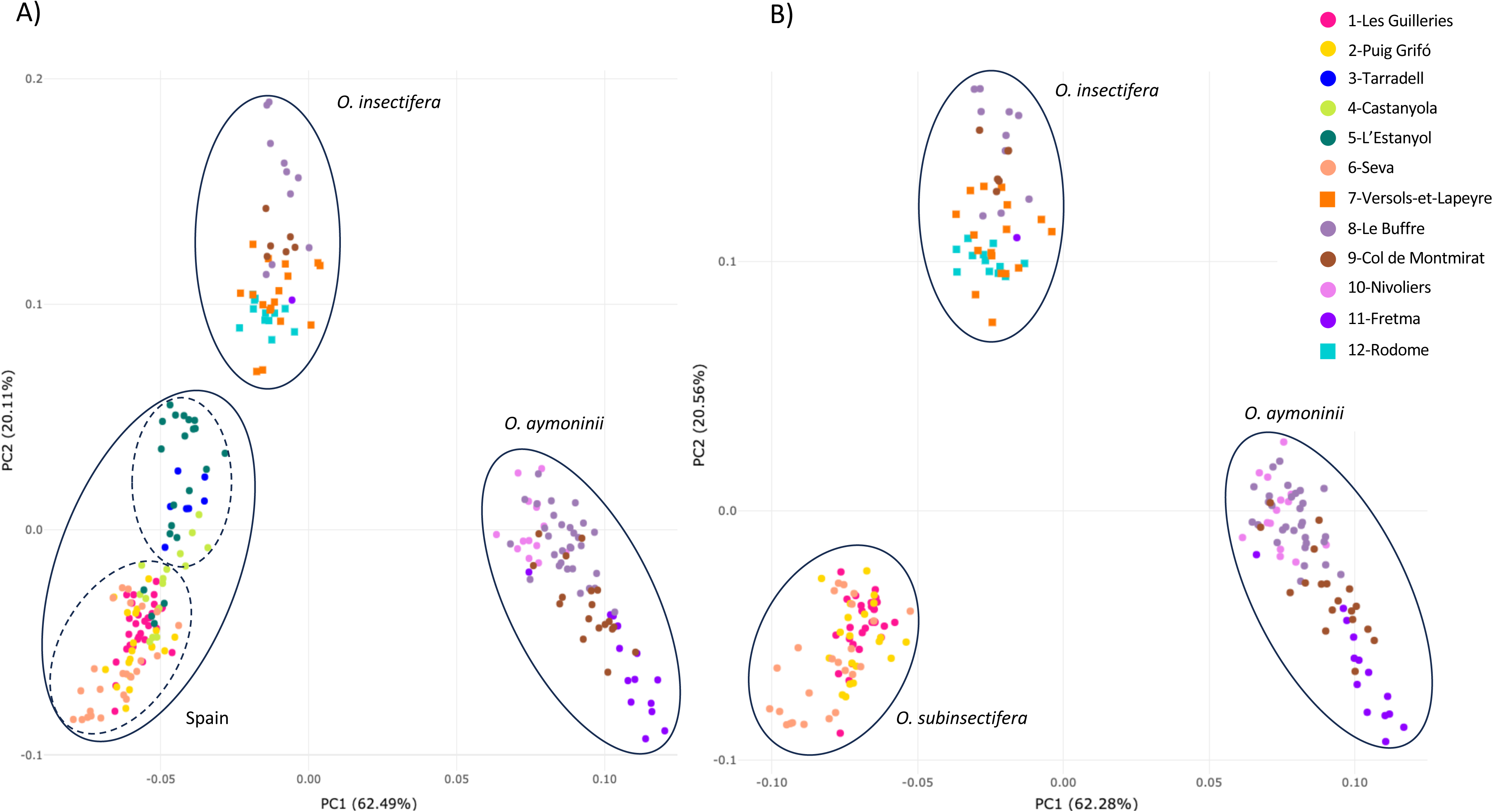
A. Principal Component Analysis (PCA) displaying the two first axes (PC1 and PC2) representing 62.49% and 20.11% of the total genetic variance. PCA was computed based on 239 individuals from 12 sites genotyped at 11,089 SNPs and colors depict sampling localities. **B** PCA displaying the two first axes (PC1 and PC2) representing 62.28% and 20.56% of the total genetic variance. PCA was computed based on 196 individuals by removing 3 sites (i.e., 3-Tarradell, 4-Castanyola and 5-Seva) and 8,611 SNPs. *O. insectifera* control sites are represented by squares, solid ellipses distinguish the three clusters and dashed ellipses characterise the subclusters.

Similarly, the clustered co-ancestry matrix obtained from FineRADstructure (Fig. 4) shows three main clusters, grouping together *O. insectifera* and *O. subinsectifera* and suggesting less similarity with *O*. *aymoninii*. A first cluster included all *O. aymoninii* individuals together. A second one grouped all *O. insectifera* from France together, regardless their site of origin. The third cluster was found to be divided into two subclusters, one consisting of intermediate individuals from the sites 3-Tarradell, 4-Castanyola and 5-L’Estanyol, and the other one consisting of *O. subinsectifera* from sites 1-Les Guilleries, 2-Puig Grifó, 6-Seva. However, four individuals from 5-L’Estanyol (*i.e*., 22-Div-025, 22-Div-026, 22-Div-027, 22-Div-044) and one individual from 4-Castanyola (22-Div-073) are clustered with *O. subinsectifera* from sites 1-Les Guilleries, 2-Puig Grifó, 6-Seva. Spanish individuals exhibit higher rates of relatedness between them among sites compared to French individuals. Indeed, sites 3-Tarradell, 4-Castanyola and 5-L’Estanyol which contain intermediate individuals display particularly high rate of relatedness. Conversely, in France, site 11-Fretma exhibit relatively high relatedness among individuals.

**Figure 4.**
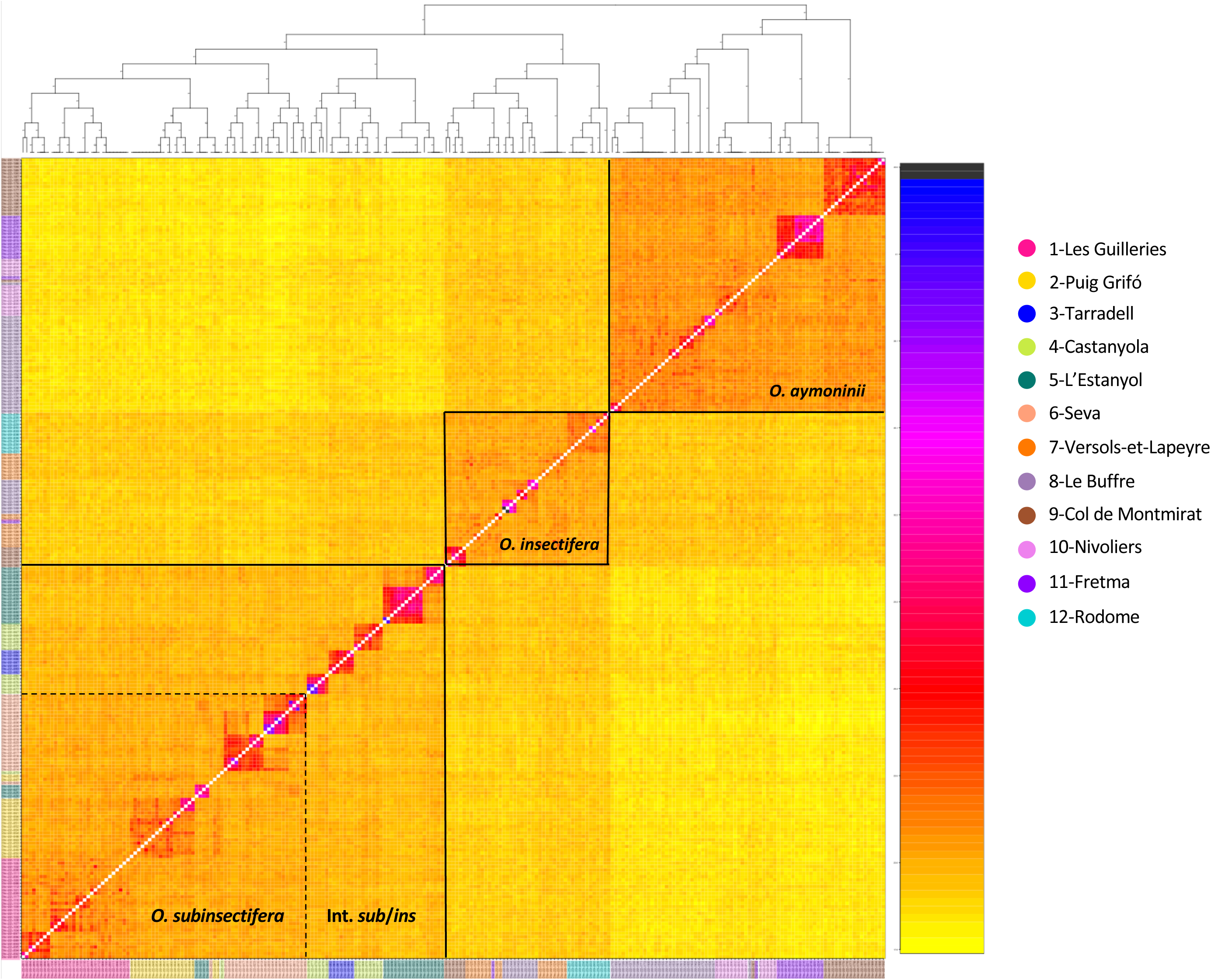
Co-ancestry matrix between each pair of individuals inferred using fineRADstructure. Each pixel depicts the magnitude of the individual co-ancestry coefficient between two individuals. Low co-ancestry coefficients values are depicted by yellow colors, whereas high values are indicated by darker colors. Different sampling localities are represented by the different colors on the side.

Furthermore, the clustering analysis carried out with ADMIXTURE suggests an optimal number of genetic clusters of *K*=8 based on the cross-entropy criterion (Supporting Information Fig. S3). We plotted the Q-matrices (that contain individual admixture coefficients) corresponding to the best replicate runs for *K*=3 (to investigate whether the program could distinguish the three taxa), *K*=4 and *K*=8. Notably, the *K*=3 graph distinguishes well *O. aymoninii*, *O. insectifera* from France, and *O. subinsectifera* from sites 1-Les Guilleries, 2-Puig Grifó, 6-Seva in light blue as well as four individuals from the site 5-L’Estanyol (22-Div-25, 22-Div-26, 22-Div-27, 22-Div-44). However, the individuals from site 3-Tarradell and the remains individuals from sites 4-Castanyola (22-Div-73, 22-Div-77, 22-Div-78, 22-Div79, 22-Div-80) and 5-L’Estanyol displayed intermediate admixture proportions (between 20 and 80%) between the two last clusters described. The pattern is overall similar at *K*=4 and *K*=8 even if they highlight particular sites like the site 11-Fretma which seems to be apart from *O. aymoninii* of the other sites or group of individuals that may be considered as relatives.

## Discussion

The population genomic approach utilised in this study, for the first time on the *Ophrys insectifera* group, confidently revealed differentiation patterns that were not discernible through morphology.

### Patterns of differentiation among the representatives of the *Ophrys insectifera* clade in Southwestern Europe

In term of morphology, our results are consistent with previous findings showing that the flowers of both endemics (*O. aymoninii* and *O. subinsectifera*) are smaller than those of the widespread *O. insectifera* (Triponez *et al*. 2013; Gervasi *et al*. 2017; Delforge 2021) . This pattern is likely to have evolved as a result of adaptation to smaller pollinator insect species: *Andrena combinata* (for *O. aymoninii*) and *Sterictiphora gastrica* (for *O. subinsectifera*) compared to the larger *Argogorytes mystaceus* (or *A. fargei*) for *O. insectifera*. In addition to dimensions, the flowers of the three taxa also show different proportions. The flowers of *O. subinsectifera* look like small flowers of *O. insectifera* with very short petals whereas the flowers of *O. aymoninii* have labellas that are relatively wider than those of *O. insectifera*, but with similar petal length. Both endemic taxa have yellow margins on their labellum and *O. aymoninii* tend to have yellowish or greenish petals, which contrast with the generally brown petals observed in the other two species. At non-floral traits, *O. insectifera* individuals tend to be taller and have more flowers but there is non-negligible overlap between species, as well as a significant amount of intra-specific variation, presumably influenced by local environmental factors, as already suggested by (Gervasi *et al*. 2017).

On the genetic data, our results clearly identified three genetic clusters whereas previous studies generally failed to distinguish the three species (Figs. 3, 4 and Supporting Information Fig. S3). Two of these clusters correspond to *O. aymoninii* and French individuals of *O. insectifera*. The two taxa are genetically distinct with no evidence of recent hybridization even at sites (e.g., 8-Le Buffre, 9-Col de Montmirat and 11-Fretma) where individuals of both species co-occur and grow at less than one meter apart. Even the three individuals that we identified as phenotypically intermediate between *O. aymoninii* and *O. insectifera* (i.e., 22-Div-146; 22-Div-152 and 22-Div-195) could be genetically assigned to *O. insectifera* for 22-Div-146 and 22-Div-195, and to *O. aymoninii* for 22-Div-152. The species barrier between *O. insectifera* and its counterpart from the Grands Causses (*O. aymoninii*) thus appears to be significant. This situation contrasts with the pattern observed on the other side of the Pyrenees. On the Spanish side, we observed a clear clustering of individuals corresponding to *O. subinsectifera*, composed of the sites 1-Les Guilleries, 2-Puig Grifó, 6-Seva and some individuals from site 4-Tarradell and 5-L’Estanyol (i.e., 22-Div-25, 22-Div-26, 22-Div-27, 22-Div-44, 22-Div-73 which are consistent with all the genetic analyses). All other Spanish individuals form a subcluster which appears as intermediate between *O. subinsectifera* and *O. insectifera*. This later subcluster is composed of all the remaining individuals from sites 4-Tarradell, 5-L’Estanyol and all the individuals from site 3-Tarradell. This suggests that the barriers to gene flow between *O. insectifera* and *O. subinsectifera* are not as strongly established as between *O. insectifera* and *O. aymoninii*. The situation becomes clearer when the individuals of those later three sites are excluded (i.e., if only sites 1-Les Guilleries, 2-Puig Grifó, 6-Seva are retained) in a way that we observe a genetic cluster formed of all *O. subinsectifera*, which is distinct from the ones formed by *O. aymoninii* and the *O. insectifera* individuals from France.

### Pervasive gene flow suspected in the Iberian Peninsula

In contrast to the observations made for *O. insectifera* and *O. aymoninii* in France, the genetic differentiation pattern suggests that at least some sites in Spain may be genetic hybrid swarms despite the individuals being clearly morphologically assignable to either *O. insectifera* or *O. subinsectifera* in most cases. The intermediate proportions of admixture shown by these individuals support this hypothesis. Notably, these admixture proportions do not allow predicting the phenotype category, whether it is *O. insectifera*, *O. subinsectifera,* or an intermediate between the two. Nevertheless, this result calls into question the existence of genetically ‘pure’ *O. insectifera* on the Iberian Peninsula. In this respect, the inclusion of an individual identified as *Ophrys insectifera* from Bercedo (Castilla y Léon), a region of Spain where *O. subinsectifera* should not be present, and genotyped using the same protocol (Gibert *et al*. 2023) also presents an intermediate status between French (at least with those from sites 7-Versols-et-Lapeyre and 12-Rodome) and Spanish individuals (Supporting Information, Fig. S4), suggesting that patterns of inter-specific gene flow may have varied from east to west of the Pyrenees, but that the existence of genetically unadmixed *O. insectifera* in Northern Spain is questionable. The possibility of inter-specific gene flow may also be supported by *G*_IS_ values that appear to be closer to panmixia expectations in Spain.

Patterns of presumed hybridization in Spain do not appear to be arranged according to geography nor ecology. All the sampling sites studied in this study are less than 10 km apart from each other and do not appear to show strong environmental differentiation. In *Ophrys*, reproductive barriers are generally thought to be pre-zygotic and maintained by specific insect pollinator species. Despite this relatively high degree of plant-insect relationship, it is not necessarily 1-to-1 (Joffard *et al*. 2019), and in some cases, secondary pollinator species may be responsible for inter-specific gene flow. To date, the sawfly *Sterictiphora gastrica* has been the only pollinator species documented for *O. subinsectifera.* Notably, on 11.05.2023, we observed and captured a small solitary bee with several pollinia attached to its head, later formally identified as *Andrena afzeliella* (by D. Genoud), pseudo-copulating on an individual of *O. subinsectifera* at site 1-Les Guilleries. Whether or not this species may be responsible for inter-specific gene flow between *Ophrys* species remains to be proven, but this observation documents a new pollinator species for the endemic *O. subinsectifera* and supports the hypothesis that secondary pollinators may play a role in the evolution of *Ophrys* (see also Claessens and Kleynen 2016 mentionning *Chalicodoma sicula* as additional pollinator).

### On the systematics and taxonomy of the *O. insectifera* clade

Our results confirm that population genomic approaches such as the one used in this study (*i.e.* nGBS) are fruitful for discriminating *Ophrys* lineages that have recently begun to diverge, and show for the first time that this is also the case for the *Ophrys insectifera* group. Even in the sampling localities where both species occur together, we could unambiguously distinguish both endemics from the widespread *Ophrys insectifera* genetically (as we could on the basis of morphology) with the notable exception of a couple Iberian individuals being intermediate between *O. insectifera* and *O. subinsectifera*. This situation confirms that pairs of closely related *Ophrys* taxa may show variable patterns of genetic differentiation between regions, as suggested by previous work (e.g. Soliva and Widmer 2003). Indeed, neutral genetic differentiation between allopatric populations of a given *Ophrys* taxon is sometimes higher than genetic differentiation between sympatric different *Ophrys* species. One hypothesis to explain this pattern is that even a small amount of pervasive inter-specific gene flow may be locally sufficient to erase the already weak level of differentiation between incipient species.

Furthermore, although the three taxa can be to some extent separated on the basis of population genomic data, we still could not reliably infer the systematic relationships between them. The widespread and generalist *O. insectifera* could intuitively be considered as occupying a basal position in the phylogenetic species tree of the group, but our results, as well as phylogenetic reconstruction rather suggest a basal position for *O. aymoninii* (Supporting Information, Fig. S5). This could support an unexpected status of relictual ancestral lineage for the micro-endemic *Ophrys aymoninii*. However, secondary contact and inter-specific gene flow after the initial divergence between *O. subinsectifera* and *O. insectifera* may also make them cluster together. Aside, gene flow between *O. aymoninii* and another species outside the *insectifera* clade could have given this pattern. This needs to be tested, but in the field, we observed putative hybrids between *O. aymoninii* and a local representative of another *Ophrys* lineage, the *O. sphegodes* group because of which introgression may also alter the position of *O. aymoninii* in the species tree.

## Conclusion

This study confirms previous research corroborating that the endemics (*O. aymoninii* and *O. subinsectifera*) can be easily distinguished from the widespread *O. insectifera* on the basis of size, proportion and colour of their flowers. Similar patterns are observed between both endemics suggesting parallel adaptation to smaller but distinct pollinators, with notably shorter labellum length and the presence yellow margins. At the genome level, this population genomic approach allowed us to differentiate for the first time both endemics from *O. insectifera* and highlight a putative hybrid swarm between *O. subinsectifera* and *O. insectifera*. This pattern, observed only in Spain could be due to recent interspecific gene flow linked to a secondary pollinator. Although further evidence is required to confirm this, this phenomenon could bias the inference of the systematic relationships between these three species by erasing the already weak phylogenetic signal.

## Acknowledgments

This work was primarily supported by an Agence Nationale pour la Recherche Jeune Chercheur Jeune Chercheuse (ANR JCJC) grant to J.B., grant number ANR-21-CE02-0022-01 and is set within the framework of the “Laboratoires d’Excellences (LABEX)” TULIP [ANR-10-LABX-41]. We thank P. Aymerich, F. Hopkins, R. Rudelle, S. Coisne, L. Barré, the Parc National des Cévennes and Takh as well as the Bertrand and Zuliani families for their support.

## Credit Statement

Salvado Pascaline (collected the data, performed the analyses and wrote the first version of the manuscript), Anaïs Gibert (collected the data, performed analyses), Bertrand Schatz (designed the study and collected the data), Lucas Vandenabeele (collected the data), Roselyne Buscail (collected the data), David Vilasís (collected the data), Philippe Feldmann (collected the data) and Joris A. M. Bertrand (designed the study, collected the data, performed the analyses and wrote the first version of the manuscript). All authors contributed substantially to the revisions.

## Data availability statement

Sequencing data have been submitted to the European Nucleotide Archive (ENA; https://www.ebi.ac.uk/ena/browser/home) under Study with primary accession n°PRJEB75177 (and secondary accession number ERA2969608277).

## Supplementary data

**Table S1.** Individual ID, Country, Population and Collector names.

**Table S2.** Mean and Standard Deviation for Floral and Whole-plant traits for the three species *O. aymoninii*, *O. insectifera*, *O. subinsectifera* and their intermediates Int. *aym*/*ins* and Int. *sub*/*ins*. All the traits are in mm, except for Plant Size, Distance to First Flower and Distance between First & Second Flower which are in cm, and the Number of Flowers & Buds.

**Table S3.** Series of *F*-statistics within individual, among individual, among populations and among Series A (between *O. aymoninii* and *O. insectifera*) and Series B (between *O. subinsectifera* and *O. insectifera*).

**Table S4.** Pairwise genetic differentiation (*G*’_ST_ values; Nei, 1987) and pairwise geographic distances (in km). All values were found to be statistically significant (*p* < 0.05).

**Figure S1.** Distribution of values for Floral and Whole-plant traits. Dashed lines represent the median value for each species and their intermediates, depicted by the different colors.

**Figure S2.** Forest plots of the different Floral and Whole-plant traits measured for each species and their intermediates. Significant *p*-values are highlighted in blue.

**Figure S3. A)** Values of the cross-entropy criterion for a number of clusters ranging from *K*=1 to 12. The optimal number of *K* was found to be 8. **B)** Barplot of ancestry coefficients obtained from ADMIXTURE for 239 individuals for *K*=3, *K*=4 and *K*=8, based on 11,089 SNPs.

**Figure S4.** Principal Component Analysis (PCA) displaying the two first axes (PC1 and PC2) representing 62.25% and 20.28% of the total genetic variance. PCA was computed on 240 individuals from the 12 sites included in this study and the site Bercedo genotyped using the same protocol (Gibert *et al*., 2023) at 11,534 SNPs and colors depict sampling localities.

**Figure S5.** Phylogenetic tree generated using Astral-III (Zhang *et al*. 2018) based on the concatenation of 4727 locus trees generated with IQ-TREE (Minh *et al*. 2020) with the best model GTR + I + G. The sites (3-Tarradell, 4-Castanyola and 5-L’Estanyol) with intermediate individuals were removed from the analysis. The three species *O. aymoninii*, *O. subinsectifera* and *O. insectifera* were found to be monophyletic with a bootstrap of 1, with the exception of two individuals 22-Div-220 and 23-Div-230, which were not phenotypically different but might be cryptically introgressed.

