## Supplementary Tables for "Contrasting patterns of differentiation among three taxa of the rapidly diversifying orchid genus *Ophrys* sect. *Insectifera* (Orchidaceae) where their range overlap"

| **Individual ID** | **Country** | **Population** | **Collectors** |
| --- | --- | --- | --- |
| Div-22-001 | Spain | 6-Seva | JB, AG, BS, RB, LV, PS, DV |
| Div-22-002 | Spain | 6-Seva | JB, AG, BS, RB, LV, PS, DV |
| Div-22-003 | Spain | 6-Seva | JB, AG, BS, RB, LV, PS, DV |
| Div-22-004 | Spain | 6-Seva | JB, AG, BS, RB, LV, PS, DV |
| Div-22-005 | Spain | 6-Seva | JB, AG, BS, RB, LV, PS, DV |
| Div-22-006 | Spain | 6-Seva | JB, AG, BS, RB, LV, PS, DV |
| Div-22-007 | Spain | 6-Seva | JB, AG, BS, RB, LV, PS, DV |
| Div-22-008 | Spain | 6-Seva | JB, AG, BS, RB, LV, PS, DV |
| Div-22-009 | Spain | 6-Seva | JB, AG, BS, RB, LV, PS, DV |
| Div-22-010 | Spain | 6-Seva | JB, AG, BS, RB, LV, PS, DV |
| Div-22-011 | Spain | 6-Seva | JB, AG, BS, RB, LV, PS, DV |
| Div-22-012 | Spain | 6-Seva | JB, AG, BS, RB, LV, PS, DV |
| Div-22-013 | Spain | 6-Seva | JB, AG, BS, RB, LV, PS, DV |
| Div-22-014 | Spain | 6-Seva | JB, AG, BS, RB, LV, PS, DV |
| Div-22-015 | Spain | 6-Seva | JB, AG, BS, RB, LV, PS, DV |
| Div-22-016 | Spain | 6-Seva | JB, AG, BS, RB, LV, PS, DV |
| Div-22-017 | Spain | 6-Seva | JB, AG, BS, RB, LV, PS, DV |
| Div-22-018 | Spain | 6-Seva | JB, AG, BS, RB, LV, PS, DV |
| Div-22-019 | Spain | 6-Seva | JB, AG, BS, RB, LV, PS, DV |
| Div-22-020 | Spain | 6-Seva | JB, AG, BS, RB, LV, PS, DV |
| Div-22-021 | Spain | 6-Seva | JB, AG, BS, RB, LV, PS, DV |
| Div-22-022 | Spain | 6-Seva | JB, AG, BS, RB, LV, PS, DV |
| Div-22-023 | Spain | 6-Seva | JB, AG, BS, RB, LV, PS, DV |
| Div-22-024 | Spain | 6-Seva | JB, AG, BS, RB, LV, PS, DV |
| Div-22-025 | Spain | 5-L'Estanyol | JB, AG, BS, RB, LV, PS, DV |
| Div-22-026 | Spain | 5-L'Estanyol | JB, AG, BS, RB, LV, PS, DV |
| Div-22-027 | Spain | 5-L'Estanyol | JB, AG, BS, RB, LV, PS, DV |
| Div-22-028 | Spain | 5-L'Estanyol | JB, AG, BS, RB, LV, PS, DV |
| Div-22-029 | Spain | 5-L'Estanyol | JB, AG, BS, RB, LV, PS, DV |
| Div-22-030 | Spain | 5-L'Estanyol | JB, AG, BS, RB, LV, PS, DV |
| Div-22-031 | Spain | 5-L'Estanyol | JB, AG, BS, RB, LV, PS, DV |
| Div-22-032 | Spain | 5-L'Estanyol | JB, AG, BS, RB, LV, PS, DV |
| Div-22-033 | Spain | 5-L'Estanyol | JB, AG, BS, RB, LV, PS, DV |
| Div-22-034 | Spain | 5-L'Estanyol | JB, AG, BS, RB, LV, PS, DV |
| Div-22-035 | Spain | 5-L'Estanyol | JB, AG, BS, RB, LV, PS, DV |
| Div-22-036 | Spain | 5-L'Estanyol | JB, AG, BS, RB, LV, PS, DV |
| Div-22-037 | Spain | 5-L'Estanyol | JB, AG, BS, RB, LV, PS, DV |
| Div-22-038 | Spain | 5-L'Estanyol | JB, AG, BS, RB, LV, PS, DV |
| Div-22-039 | Spain | 5-L'Estanyol | JB, AG, BS, RB, LV, PS, DV |
| Div-22-040 | Spain | 5-L'Estanyol | JB, AG, BS, RB, LV, PS, DV |
| Div-22-041 | Spain | 5-L'Estanyol | JB, AG, BS, RB, LV, PS, DV |
| Div-22-042 | Spain | 5-L'Estanyol | JB, AG, BS, RB, LV, PS, DV |
| Div-22-043 | Spain | 5-L'Estanyol | JB, AG, BS, RB, LV, PS, DV |
| Div-22-044 | Spain | 5-L'Estanyol | JB, AG, BS, RB, LV, PS, DV |
| Div-22-045 | Spain | 5-L'Estanyol | JB, AG, BS, RB, LV, PS, DV |
| Div-22-046 | Spain | 2-Puig Grifó | JB, AG, BS, RB, LV, PS, DV |
| Div-22-047 | Spain | 2-Puig Grifó | JB, AG, BS, RB, LV, PS, DV |
| Div-22-048 | Spain | 2-Puig Grifó | JB, AG, BS, RB, LV, PS, DV |
| Div-22-049 | Spain | 2-Puig Grifó | JB, AG, BS, RB, LV, PS, DV |
| Div-22-050 | Spain | 2-Puig Grifó | JB, AG, BS, RB, LV, PS, DV |
| Div-22-051 | Spain | 2-Puig Grifó | JB, AG, BS, RB, LV, PS, DV |
| Div-22-052 | Spain | 2-Puig Grifó | JB, AG, BS, RB, LV, PS, DV |
| Div-22-053 | Spain | 2-Puig Grifó | JB, AG, BS, RB, LV, PS, DV |
| Div-22-054 | Spain | 2-Puig Grifó | JB, AG, BS, RB, LV, PS, DV |
| Div-22-055 | Spain | 2-Puig Grifó | JB, AG, BS, RB, LV, PS, DV |
| Div-22-056 | Spain | 2-Puig Grifó | JB, AG, BS, RB, LV, PS, DV |
| Div-22-057 | Spain | 2-Puig Grifó | JB, AG, BS, RB, LV, PS, DV |
| Div-22-058 | Spain | 2-Puig Grifó | JB, AG, BS, RB, LV, PS, DV |
| Div-22-059 | Spain | 2-Puig Grifó | JB, AG, BS, RB, LV, PS, DV |
| Div-22-060 | Spain | 2-Puig Grifó | JB, AG, BS, RB, LV, PS, DV |
| Div-22-061 | Spain | 2-Puig Grifó | JB, AG, BS, RB, LV, PS, DV |
| Div-22-062 | Spain | 2-Puig Grifó | JB, AG, BS, RB, LV, PS, DV |
| Div-22-063 | Spain | 2-Puig Grifó | JB, AG, BS, RB, LV, PS, DV |
| Div-22-064 | Spain | 2-Puig Grifó | JB, AG, BS, RB, LV, PS, DV |
| Div-22-065 | Spain | 2-Puig Grifó | JB, AG, BS, RB, LV, PS, DV |
| Div-22-066 | Spain | 4-Castanyola | JB, AG, BS, RB, LV, PS, DV |
| Div-22-067 | Spain | 4-Castanyola | JB, AG, BS, RB, LV, PS, DV |
| Div-22-068 | Spain | 4-Castanyola | JB, AG, BS, RB, LV, PS, DV |
| Div-22-069 | Spain | 4-Castanyola | JB, AG, BS, RB, LV, PS, DV |
| Div-22-070 | Spain | 4-Castanyola | JB, AG, BS, RB, LV, PS, DV |
| Div-22-071 | Spain | 4-Castanyola | JB, AG, BS, RB, LV, PS, DV |
| Div-22-072 | Spain | 4-Castanyola | JB, AG, BS, RB, LV, PS, DV |
| Div-22-073 | Spain | 4-Castanyola | JB, AG, BS, RB, LV, PS, DV |
| Div-22-074 | Spain | 4-Castanyola | JB, AG, BS, RB, LV, PS, DV |
| Div-22-075 | Spain | 4-Castanyola | JB, AG, BS, RB, LV, PS, DV |
| Div-22-076 | Spain | 4-Castanyola | JB, AG, BS, RB, LV, PS, DV |
| Div-22-077 | Spain | 4-Castanyola | JB, AG, BS, RB, LV, PS, DV |
| Div-22-078 | Spain | 4-Castanyola | JB, AG, BS, RB, LV, PS, DV |
| Div-22-079 | Spain | 4-Castanyola | JB, AG, BS, RB, LV, PS, DV |
| Div-22-080 | Spain | 4-Castanyola | JB, AG, BS, RB, LV, PS, DV |
| Div-22-081 | Spain | 3-Tarradell | JB, AG, BS, RB, LV, PS, DV |
| Div-22-082 | Spain | 3-Tarradell | JB, AG, BS, RB, LV, PS, DV |
| Div-22-083 | Spain | 3-Tarradell | JB, AG, BS, RB, LV, PS, DV |
| Div-22-084 | Spain | 3-Tarradell | JB, AG, BS, RB, LV, PS, DV |
| Div-22-085 | Spain | 3-Tarradell | JB, AG, BS, RB, LV, PS, DV |
| Div-22-086 | Spain | 3-Tarradell | JB, AG, BS, RB, LV, PS, DV |
| Div-22-087 | Spain | 3-Tarradell | JB, AG, BS, RB, LV, PS, DV |
| Div-22-088 | Spain | 1-Les Guilleries | JB, AG, BS, RB, LV, PS, DV |
| Div-22-089 | Spain | 1-Les Guilleries | JB, AG, BS, RB, LV, PS, DV |
| Div-22-090 | Spain | 1-Les Guilleries | JB, AG, BS, RB, LV, PS, DV |
| Div-22-091 | Spain | 1-Les Guilleries | JB, AG, BS, RB, LV, PS, DV |
| Div-22-092 | Spain | 1-Les Guilleries | JB, AG, BS, RB, LV, PS, DV |
| Div-22-093 | Spain | 1-Les Guilleries | JB, AG, BS, RB, LV, PS, DV |
| Div-22-094 | Spain | 1-Les Guilleries | JB, AG, BS, RB, LV, PS, DV |
| Div-22-095 | Spain | 1-Les Guilleries | JB, AG, BS, RB, LV, PS, DV |
| Div-22-096 | Spain | 1-Les Guilleries | JB, AG, BS, RB, LV, PS, DV |
| Div-22-097 | Spain | 1-Les Guilleries | JB, AG, BS, RB, LV, PS, DV |
| Div-22-098 | Spain | 1-Les Guilleries | JB, AG, BS, RB, LV, PS, DV |
| Div-22-099 | Spain | 1-Les Guilleries | JB, AG, BS, RB, LV, PS, DV |
| Div-22-100 | Spain | 1-Les Guilleries | JB, AG, BS, RB, LV, PS, DV |
| Div-22-101 | Spain | 1-Les Guilleries | JB, AG, BS, RB, LV, PS, DV |
| Div-22-102 | Spain | 1-Les Guilleries | JB, AG, BS, RB, LV, PS, DV |
| Div-22-103 | Spain | 1-Les Guilleries | JB, AG, BS, RB, LV, PS, DV |
| Div-22-104 | Spain | 1-Les Guilleries | JB, AG, BS, RB, LV, PS, DV |
| Div-22-105 | Spain | 1-Les Guilleries | JB, AG, BS, RB, LV, PS, DV |
| Div-22-106 | Spain | 1-Les Guilleries | JB, AG, BS, RB, LV, PS, DV |
| Div-22-107 | Spain | 1-Les Guilleries | JB, AG, BS, RB, LV, PS, DV |
| Div-22-108 | Spain | 1-Les Guilleries | JB, AG, BS, RB, LV, PS, DV |
| Div-22-109 | Spain | 1-Les Guilleries | JB, AG, BS, RB, LV, PS, DV |
| Div-22-110 | Spain | 1-Les Guilleries | JB, AG, BS, RB, LV, PS, DV |
| Div-22-111 | Spain | 1-Les Guilleries | JB, AG, BS, RB, LV, PS, DV |
| Div-22-112 | Spain | 1-Les Guilleries | JB, AG, BS, RB, LV, PS, DV |
| Div-22-113 | Spain | 1-Les Guilleries | JB, AG, BS, RB, LV, PS, DV |
| Div-22-114 | Spain | 1-Les Guilleries | JB, AG, BS, RB, LV, PS, DV |
| Div-22-115 | Spain | 1-Les Guilleries | JB, AG, BS, RB, LV, PS, DV |
| Div-22-116 | Spain | 1-Les Guilleries | JB, AG, BS, RB, LV, PS, DV |
| Div-22-117 | Spain | 1-Les Guilleries | JB, AG, BS, RB, LV, PS, DV |
| Div-22-118 | France | 7-Versols-et-Lapeyre | JB, AG |
| Div-22-119 | France | 7-Versols-et-Lapeyre | JB, AG |
| Div-22-120 | France | 7-Versols-et-Lapeyre | JB, AG |
| Div-22-121 | France | 7-Versols-et-Lapeyre | JB, AG |
| Div-22-122 | France | 7-Versols-et-Lapeyre | JB, AG |
| Div-22-123 | France | 7-Versols-et-Lapeyre | JB, AG |
| Div-22-124 | France | 7-Versols-et-Lapeyre | JB, AG |
| Div-22-125 | France | 7-Versols-et-Lapeyre | JB, AG |
| Div-22-126 | France | 7-Versols-et-Lapeyre | JB, AG |
| Div-22-127 | France | 7-Versols-et-Lapeyre | JB, AG |
| Div-22-128 | France | 7-Versols-et-Lapeyre | JB, AG |
| Div-22-129 | France | 7-Versols-et-Lapeyre | JB, AG |
| Div-22-130 | France | 7-Versols-et-Lapeyre | JB, AG |
| Div-22-131 | France | 7-Versols-et-Lapeyre | JB, AG |
| Div-22-132 | France | 7-Versols-et-Lapeyre | JB, AG |
| Div-22-133 | France | 7-Versols-et-Lapeyre | JB, AG |
| Div-22-134 | France | 7-Versols-et-Lapeyre | JB, AG |
| Div-22-135 | France | 8-Le Buffre | JB, AG, BS, RB, LV, PS |
| Div-22-136 | France | 8-Le Buffre | JB, AG, BS, RB, LV, PS |
| Div-22-137 | France | 8-Le Buffre | JB, AG, BS, RB, LV, PS |
| Div-22-138 | France | 8-Le Buffre | JB, AG, BS, RB, LV, PS |
| Div-22-139 | France | 8-Le Buffre | JB, AG, BS, RB, LV, PS |
| Div-22-140 | France | 8-Le Buffre | JB, AG, BS, RB, LV, PS |
| Div-22-141 | France | 8-Le Buffre | JB, AG, BS, RB, LV, PS |
| Div-22-142 | France | 8-Le Buffre | JB, AG, BS, RB, LV, PS |
| Div-22-143 | France | 8-Le Buffre | JB, AG, BS, RB, LV, PS |
| Div-22-144 | France | 8-Le Buffre | JB, AG, BS, RB, LV, PS |
| Div-22-145 | France | 8-Le Buffre | JB, AG, BS, RB, LV, PS |
| Div-22-146 | France | 8-Le Buffre | JB, AG, BS, RB, LV, PS |
| Div-22-147 | France | 8-Le Buffre | JB, AG, BS, RB, LV, PS |
| Div-22-148 | France | 8-Le Buffre | JB, AG, BS, RB, LV, PS |
| Div-22-149 | France | 8-Le Buffre | JB, AG, BS, RB, LV, PS |
| Div-22-150 | France | 8-Le Buffre | JB, AG, BS, RB, LV, PS |
| Div-22-151 | France | 8-Le Buffre | JB, AG, BS, RB, LV, PS |
| Div-22-152 | France | 8-Le Buffre | JB, AG, BS, RB, LV, PS |
| Div-22-153 | France | 8-Le Buffre | JB, AG, BS, RB, LV, PS |
| Div-22-154 | France | 8-Le Buffre | JB, AG, BS, RB, LV, PS |
| Div-22-155 | France | 8-Le Buffre | JB, AG, BS, RB, LV, PS |
| Div-22-156 | France | 8-Le Buffre | JB, AG, BS, RB, LV, PS |
| Div-22-157 | France | 8-Le Buffre | JB, AG, BS, RB, LV, PS |
| Div-22-158 | France | 8-Le Buffre | JB, AG, BS, RB, LV, PS |
| Div-22-159 | France | 8-Le Buffre | JB, AG, BS, RB, LV, PS |
| Div-22-160 | France | 8-Le Buffre | JB, AG, BS, RB, LV, PS |
| Div-22-161 | France | 8-Le Buffre | JB, AG, BS, RB, LV, PS |
| Div-22-162 | France | 8-Le Buffre | JB, AG, BS, RB, LV, PS |
| Div-22-163 | France | 8-Le Buffre | JB, AG, BS, RB, LV, PS |
| Div-22-164 | France | 8-Le Buffre | JB, AG, BS, RB, LV, PS |
| Div-22-165 | France | 8-Le Buffre | JB, AG, BS, RB, LV, PS |
| Div-22-166 | France | 8-Le Buffre | JB, AG, BS, RB, LV, PS |
| Div-22-167 | France | 8-Le Buffre | JB, AG, BS, RB, LV, PS |
| Div-22-168 | France | 8-Le Buffre | JB, AG, BS, RB, LV, PS |
| Div-22-169 | France | 8-Le Buffre | JB, AG, BS, RB, LV, PS |
| Div-22-170 | France | 8-Le Buffre | JB, AG, BS, RB, LV, PS |
| Div-22-171 | France | 8-Le Buffre | JB, AG, BS, RB, LV, PS |
| Div-22-172 | France | 8-Le Buffre | JB, AG, BS, RB, LV, PS |
| Div-22-173 | France | 8-Le Buffre | JB, AG, BS, RB, LV, PS |
| Div-22-174 | France | 8-Le Buffre | JB, AG, BS, RB, LV, PS |
| Div-22-175 | France | 9-Col de Montmirat | JB, AG, BS, RB, LV, PS |
| Div-22-176 | France | 9-Col de Montmirat | JB, AG, BS, RB, LV, PS |
| Div-22-177 | France | 9-Col de Montmirat | JB, AG, BS, RB, LV, PS |
| Div-22-178 | France | 9-Col de Montmirat | JB, AG, BS, RB, LV, PS |
| Div-22-179 | France | 9-Col de Montmirat | JB, AG, BS, RB, LV, PS |
| Div-22-180 | France | 9-Col de Montmirat | JB, AG, BS, RB, LV, PS |
| Div-22-181 | France | 9-Col de Montmirat | JB, AG, BS, RB, LV, PS |
| Div-22-182 | France | 9-Col de Montmirat | JB, AG, BS, RB, LV, PS |
| Div-22-183 | France | 9-Col de Montmirat | JB, AG, BS, RB, LV, PS |
| Div-22-184 | France | 9-Col de Montmirat | JB, AG, BS, RB, LV, PS |
| Div-22-185 | France | 9-Col de Montmirat | JB, AG, BS, RB, LV, PS |
| Div-22-186 | France | 9-Col de Montmirat | JB, AG, BS, RB, LV, PS |
| Div-22-187 | France | 9-Col de Montmirat | JB, AG, BS, RB, LV, PS |
| Div-22-188 | France | 9-Col de Montmirat | JB, AG, BS, RB, LV, PS |
| Div-22-189 | France | 9-Col de Montmirat | JB, AG, BS, RB, LV, PS |
| Div-22-190 | France | 9-Col de Montmirat | JB, AG, BS, RB, LV, PS |
| Div-22-191 | France | 9-Col de Montmirat | JB, AG, BS, RB, LV, PS |
| Div-22-192 | France | 9-Col de Montmirat | JB, AG, BS, RB, LV, PS |
| Div-22-193 | France | 9-Col de Montmirat | JB, AG, BS, RB, LV, PS |
| Div-22-194 | France | 9-Col de Montmirat | JB, AG, BS, RB, LV, PS |
| Div-22-195 | France | 9-Col de Montmirat | JB, AG, BS, RB, LV, PS |
| Div-22-196 | France | 9-Col de Montmirat | JB, AG, BS, RB, LV, PS |
| Div-22-197 | France | 9-Col de Montmirat | JB, AG, BS, RB, LV, PS |
| Div-22-198 | France | 9-Col de Montmirat | JB, AG, BS, RB, LV, PS |
| Div-22-199 | France | 10-Nivoliers | JB, AG, BS, RB, LV, PS |
| Div-22-200 | France | 10-Nivoliers | JB, AG, BS, RB, LV, PS |
| Div-22-201 | France | 10-Nivoliers | JB, AG, BS, RB, LV, PS |
| Div-22-202 | France | 10-Nivoliers | JB, AG, BS, RB, LV, PS |
| Div-22-204 | France | 10-Nivoliers | JB, AG, BS, RB, LV, PS |
| Div-22-205 | France | 10-Nivoliers | JB, AG, BS, RB, LV, PS |
| Div-22-206 | France | 10-Nivoliers | JB, AG, BS, RB, LV, PS |
| Div-22-207 | France | 10-Nivoliers | JB, AG, BS, RB, LV, PS |
| Div-22-208 | France | 10-Nivoliers | JB, AG, BS, RB, LV, PS |
| Div-22-209 | France | 10-Nivoliers | JB, AG, BS, RB, LV, PS |
| Div-22-210 | France | 10-Nivoliers | JB, AG, BS, RB, LV, PS |
| Div-22-211 | France | 10-Nivoliers | JB, AG, BS, RB, LV, PS |
| Div-22-212 | France | 10-Nivoliers | JB, AG, BS, RB, LV, PS |
| Div-22-213 | France | 10-Nivoliers | JB, AG, BS, RB, LV, PS |
| Div-22-214 | France | 11-Fretma | JB, AG, BS, RB, LV, PS |
| Div-22-215 | France | 11-Fretma | JB, AG, BS, RB, LV, PS |
| Div-22-216 | France | 11-Fretma | JB, AG, BS, RB, LV, PS |
| Div-22-217 | France | 11-Fretma | JB, AG, BS, RB, LV, PS |
| Div-22-218 | France | 11-Fretma | JB, AG, BS, RB, LV, PS |
| Div-22-219 | France | 11-Fretma | JB, AG, BS, RB, LV, PS |
| Div-22-220 | France | 11-Fretma | JB, AG, BS, RB, LV, PS |
| Div-22-221 | France | 11-Fretma | JB, AG, BS, RB, LV, PS |
| Div-22-222 | France | 11-Fretma | JB, AG, BS, RB, LV, PS |
| Div-22-223 | France | 11-Fretma | JB, AG, BS, RB, LV, PS |
| Div-22-224 | France | 11-Fretma | JB, AG, BS, RB, LV, PS |
| Div-22-225 | France | 11-Fretma | JB, AG, BS, RB, LV, PS |
| Div-22-226 | France | 11-Fretma | JB, AG, BS, RB, LV, PS |
| Div-22-227 | France | 11-Fretma | JB, AG, BS, RB, LV, PS |
| Div-22-228 | France | 11-Fretma | JB, AG, BS, RB, LV, PS |
| Div-23-229 | France | 12-Rodome | JB, AG, LV, PS |
| Div-23-230 | France | 12-Rodome | JB, AG, LV, PS |
| Div-23-231 | France | 12-Rodome | JB, AG, LV, PS |
| Div-23-232 | France | 12-Rodome | JB, AG, LV, PS |
| Div-23-233 | France | 12-Rodome | JB, AG, LV, PS |
| Div-23-234 | France | 12-Rodome | JB, AG, LV, PS |
| Div-23-235 | France | 12-Rodome | JB, AG, LV, PS |
| Div-23-236 | France | 12-Rodome | JB, AG, LV, PS |
| Div-23-237 | France | 12-Rodome | JB, AG, LV, PS |
| Div-23-238 | France | 12-Rodome | JB, AG, LV, PS |
| Div-23-239 | France | 12-Rodome | JB, AG, LV, PS |
| Div-23-240 | France | 12-Rodome | JB, AG, LV, PS |

Collectors: Joris Bertrand (JB), Anaïs Gibert (AG), Bertrand Schatz (BS), Roselyne Buscail (RB), Lucas Vandenabeele (LV), Pascaline Salvado (PS), David Vilasís (DV).

**Table S1.** Individual ID, Country, Population and Collector names.

| **Traits** |  | **O. aymoninii** | **Int. aym/ins** | **O. insectifera** | **Int. sub/ins** | **O. subinsectifera** |
| --- | --- | --- | --- | --- | --- | --- |
| **Labellum Length** | Mean | 9.36840 | 11.22000 | 12.96294 | 12.41786 | 10.59531 |
|  | Standard Deviation | 1.170686 | 2.008382 | 1.741534 | 1.008969 | 1.332933 |
| **Labellum Width** | Mean | 6.933200 | 5.700000 | 5.709440 | 5.996429 | 5.418115 |
|  | Standard Deviation | 1.1028417 | 2.0129829 | 1.2054504 | 0.8573269 | 1.0122579 |
| **Ratio Labellum** | Mean | 1.374591 | 2.083597 | 2.373345 | 2.103101 | 2.012991 |
|  | Standard Deviation | 0.2187878 | 0.5326773 | 0.6089824 | 0.2994695 | 0.4051127 |
| **Sepal Length** | Mean | 5.035333 | 5.406667 | 5.946760 | 6.335714 | 5.685250 |
|  | Standard Deviation | 0.7677967 | 0.6503332 | 0.8334926 | 0.7584672 | 1.0526463 |
| **Sepal Width** | Mean | 2.721067 | 2.556667 | 2.736400 | 2.480714 | 2.342596 |
|  | Standard Deviation | 0.4384203 | 0.1450287 | 0.4346254 | 0.3982165 | 0.5667435 |
| **Petal Length** | Mean | 3.517200 | 3.733333 | 3.499260 | 4.127857 | 2.656533 |
|  | Standard Deviation | 0.6950414 | 0.7487545 | 1.1185874 | 0.6699274 | 0.8917885 |
| **Plant Size** | Mean | 19.71600 | 28.63333 | 25.37400 | 23.85714 | 19.39187 |
|  | Standard Deviation | 6.291351 | 3.992910 | 5.928582 | 5.596526 | 6.373750 |
| **Stem Diameter** | Mean | 2.211867 | 2.366667 | 2.042400 | 2.527857 | 2.429421 |
|  | Standard Deviation | 0.6409615 | 0.4974267 | 0.7188328 | 0.6963551 | 0.9581294 |
| **Number of Flowers & Buds** | Mean | 6.213333 | 7.666667 | 7.320000 | 6.928571 | 6.958333 |
|  | Standard Deviation | 2.213554 | 1.154701 | 2.151791 | 2.129077 | 2.529475 |
| **Distance to First Flower** | Mean | 12.41867 | 17.63333 | 16.76200 | 15.05714 | 12.61354 |
|  | Standard Deviation | 3.506199 | 1.011599 | 4.926061 | 3.360795 | 4.178205 |
| **Distance between First & Second Flower** | Mean | 2.154667 | 2.266667 | 2.906122 | 3.818182 | 2.579268 |
|  | Standard Deviation | 0.6018650 | 1.1590226 | 0.8685256 | 1.8269199 | 1.2867547 |

**Table S2.** Mean and Standard Deviation for Floral and Whole-plant traits for the three species *O. aymoninii*, *O. insectifera*, *O. subinsectifera* and their intermediates Int. *aym*/*ins* and Int. *sub*/*ins.* All the traits are in mm, except for Plant Size, Distance to First Flower and Distance between First & Second Flower which are in cm, and the Number of Flowers & Buds.

**Series A : *O. aymoninii* vs *O. insectifera***

| Source of variation | %var | F-stat | F-value | Sdt Dev | c.i.2.5% | | c.i.97.5% | | P-value | F'-value |
| --- | --- | --- | --- | --- | --- | --- | --- | --- | --- | --- |
| Within individual | 0.785 | *F*_IT_ | 0.215 | 0.003 | 0.209 | 0.22 | |  | |  |
| Among individual | 0.084 | *F*_IS_ | 0.096 | 0.002 | 0.092 | 0.101 | | 0.001 | |  |
| Among population | 0.059 | *F*_SC_ | 0.063 | 0.001 | 0.062 | 0.064 | | 0.001 | | 0.085 |
| Among Series A | 0.072 | *F*_CT_ | 0.072 | 0.002 | 0.068 | 0.077 | | 0.001 | | 0.099 |

**Series B : *O. subinsectifera* vs *O. insectifera***

| Source of variation | %var | F-stat | F-value | Sdt Dev | c.i.2.5% | | c.i.97.5% | | P-value | F'-value |
| --- | --- | --- | --- | --- | --- | --- | --- | --- | --- | --- |
| Within individual | 0.795 | *F*_IT_ | 0.205 | 0.003 | 0.2 | 0.211 | |  | |  |
| Among individual | 0.085 | *F*_IS_ | 0.096 | 0.002 | 0.092 | 0.101 | | 0.001 | |  |
| Among population | 0.083 | *F*_SC_ | 0.086 | 0.001 | 0.084 | 0.088 | | 0.001 | | 0.115 |
| Among Series B | 0.038 | *F*_CT_ | 0.038 | 0.001 | 0.036 | 0.041 | | 0.001 | | 0.053 |

**Table S3.** Series of *F*-statistics within individual, among individual, among population and among Series A (between *O. aymoninii* and *O. insectifera*) and Series B (between *O. subinsectifera* and *O. insectifera*).

| **G'st(Nei)/**  **Distance** | **1-Les Guilleries** | **2- Puig Grifó** | **3-Tarradell** | **4-Castanyola** | **5-L’Estanyol** | **6-Seva** | **7-Versols-et-Lapeyre** | **8-Le Buffre** | **9-Col de Montmirat** | **10-Nivoliers** | **11-Fretma** | **12-Rodome** |
| --- | --- | --- | --- | --- | --- | --- | --- | --- | --- | --- | --- | --- |
| **1-Les Guilleries** |  | 4.1 | 4.76 | 7.35 | 9.28 | 8.37 | 222.85 | 281.23 | 298.22 | 278.37 | 278 | 104 |
| **2-Puig Grifó** | 0.041 |  | 2.65 | 3.73 | 5.18 | 4.27 | 232.25 | 284.83 | 301.86 | 282 | 281.65 | 106.2 |
| **3-Tarradell** | 0.102 | 0.115 |  | 3 | 6.01 | 5.17 | 230.66 | 283.45 | 300.53 | 280.66 | 280.34 | 103.8 |
| **4-Castanyola** | 0.068 | 0.071 | 0.121 |  | 3.47 | 2.84 | 233.29 | 286.2 | 303.31 | 283.43 | 283.12 | 105.7 |
| **5-L’Estanyol** | 0.06 | 0.068 | 0.127 | 0.083 |  | 0.91 | 236.64 | 289.46 | 306.54 | 286.67 | 286.35 | 109.1 |
| **6-Seva** | 0.055 | 0.067 | 0.134 | 0.088 | 0.072 |  | 235.83 | 288.61 | 305.69 | 285.82 | 285.49 | 108.4 |
| **7-Versols-et-Lapeyre** | 0.066 | 0.076 | 0.103 | 0.083 | 0.069 | 0.083 |  | 58.14 | 76.16 | 57.46 | 58.65 | 142 |
| **8-Le Buffre** | 0.122 | 0.132 | 0.157 | 0.135 | 0.121 | 0.142 | 0.084 |  | 18.03 | 5.31 | 8.31 | 195.5 |
| **9-Col de Montmirat** | 0.153 | 0.163 | 0.193 | 0.17 | 0.158 | 0.178 | 0.123 | 0.053 |  | 19.9 | 20.31 | 217.4 |
| **10-Nivoliers** | 0.114 | 0.128 | 0.155 | 0.132 | 0.12 | 0.137 | 0.079 | 0.019 | 0.054 |  | 3.2 | 195.3 |
| **11-Fretma** | 0.194 | 0.203 | 0.239 | 0.21 | 0.198 | 0.217 | 0.159 | 0.087 | 0.128 | 0.093 |  | 193.9 |
| **12-Rodome** | 0.069 | 0.077 | 0.111 | 0.087 | 0.078 | 0.09 | 0.027 | 0.096 | 0.13 | 0.087 | 0.167 |  |

**Table S4.** Pairwise genetic differentiation (*G*’_ST_ values; Nei, 1987) and pairwise geographic distances (in km). All values were found to be statistically significant (*p* < 0.05).
