## Supplementary Figures for "Contrasting patterns of differentiation among three taxa of the rapidly diversifying orchid genus *Ophrys* sect. *Insectifera* (Orchidaceae) where their range overlap"

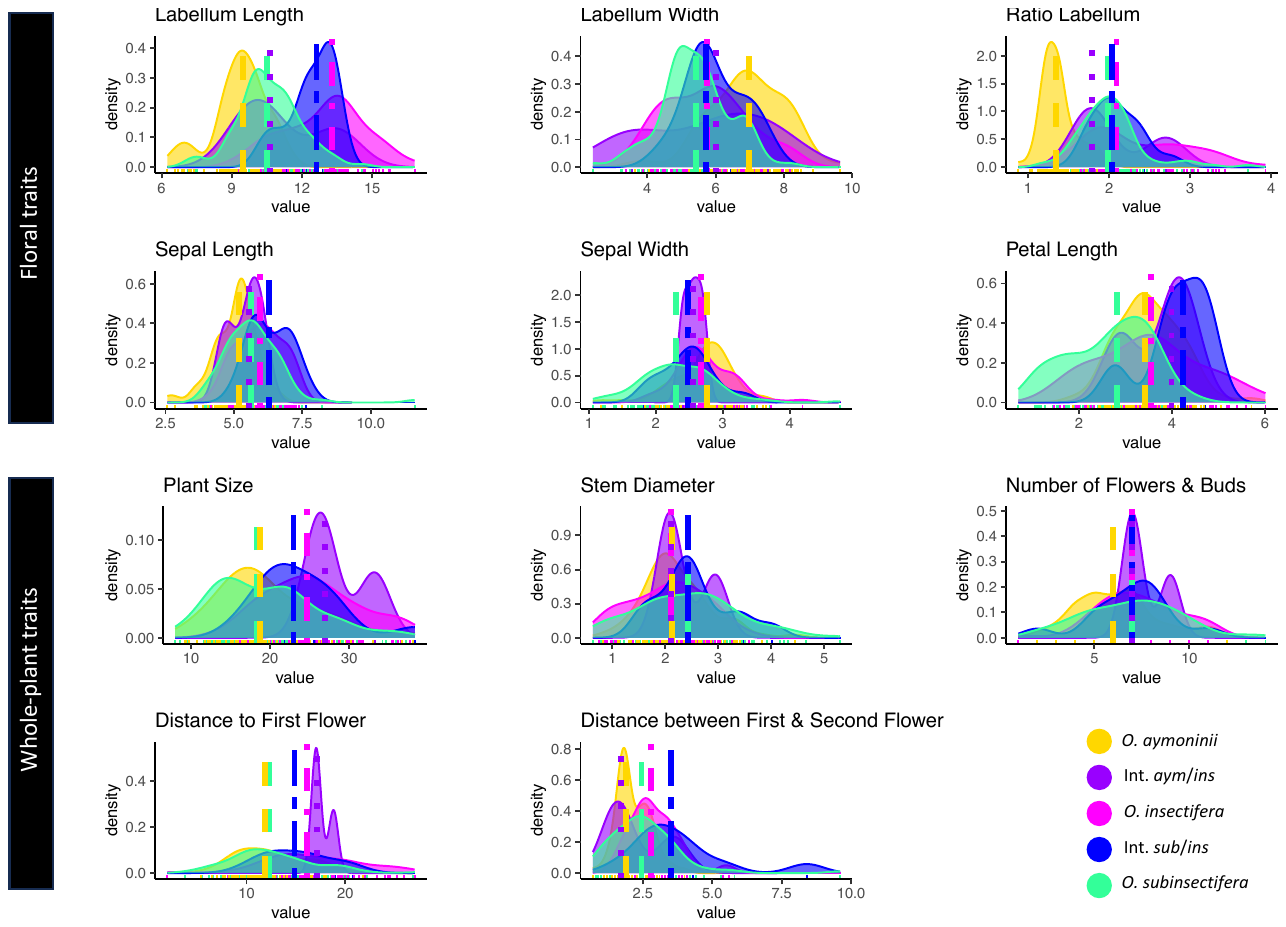
**Figure S1.** Distribution of values for Floral and Whole-plant traits. Dashed lines represent the median value for each species and their intermediates, depicted by the different colors.

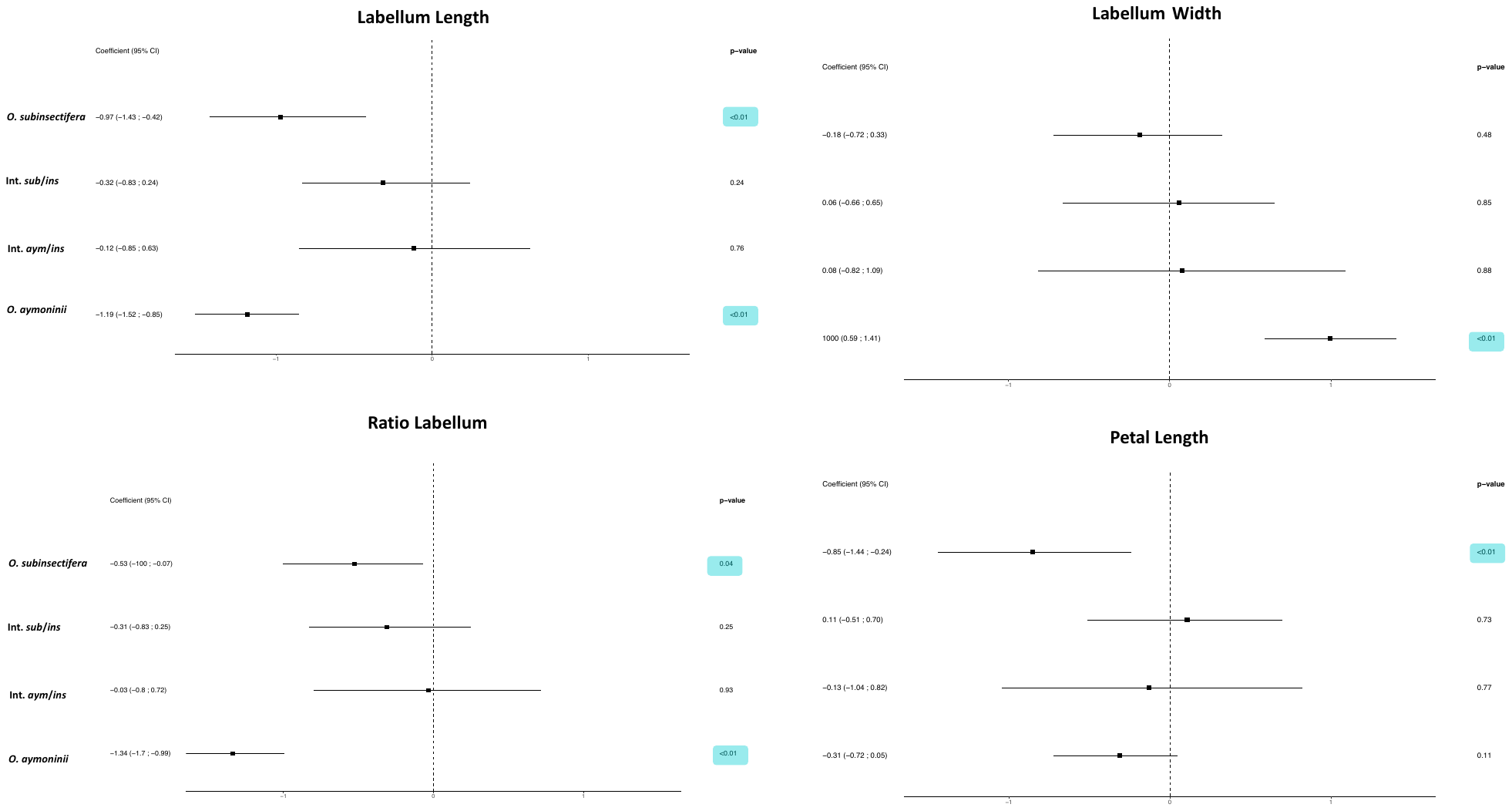

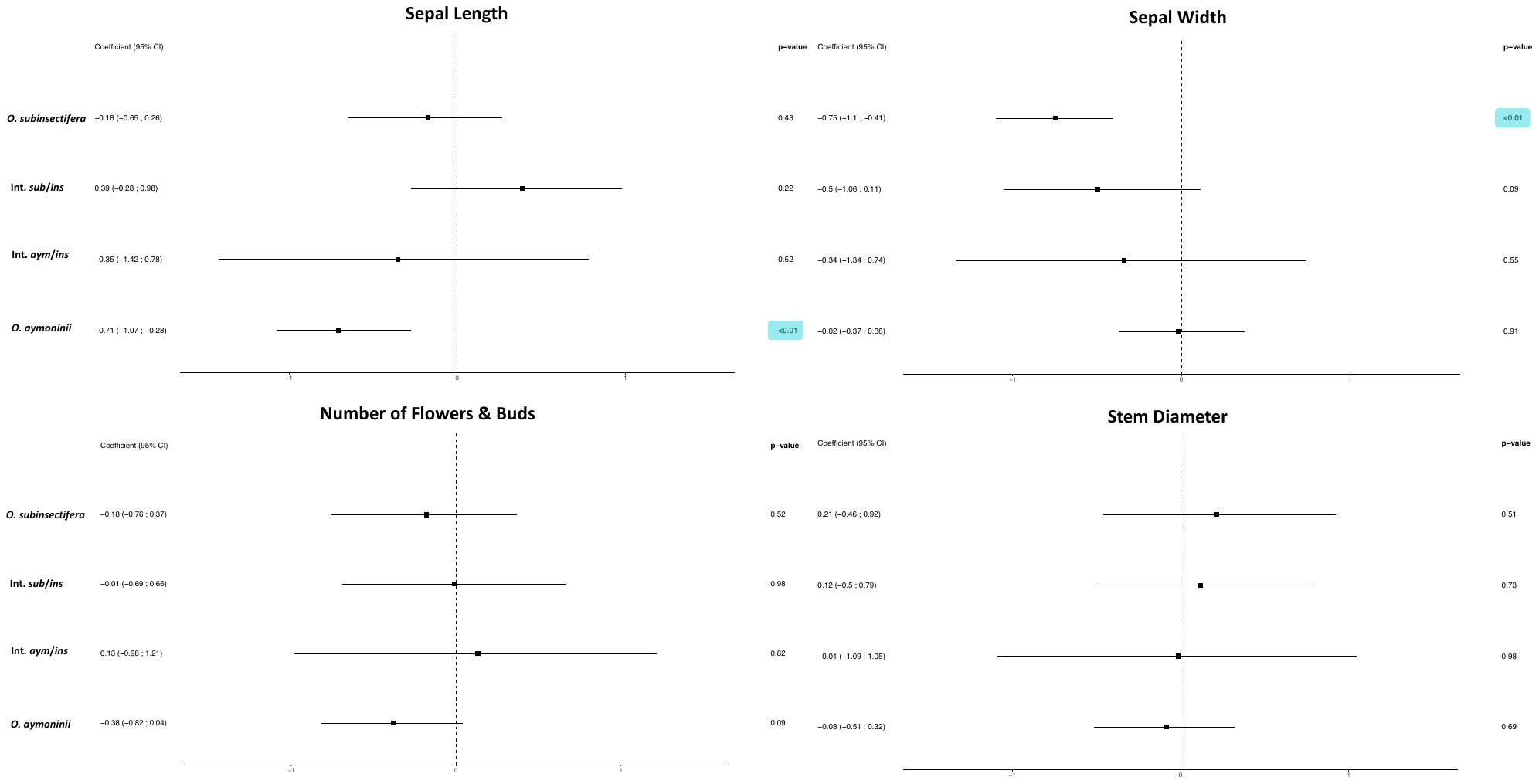

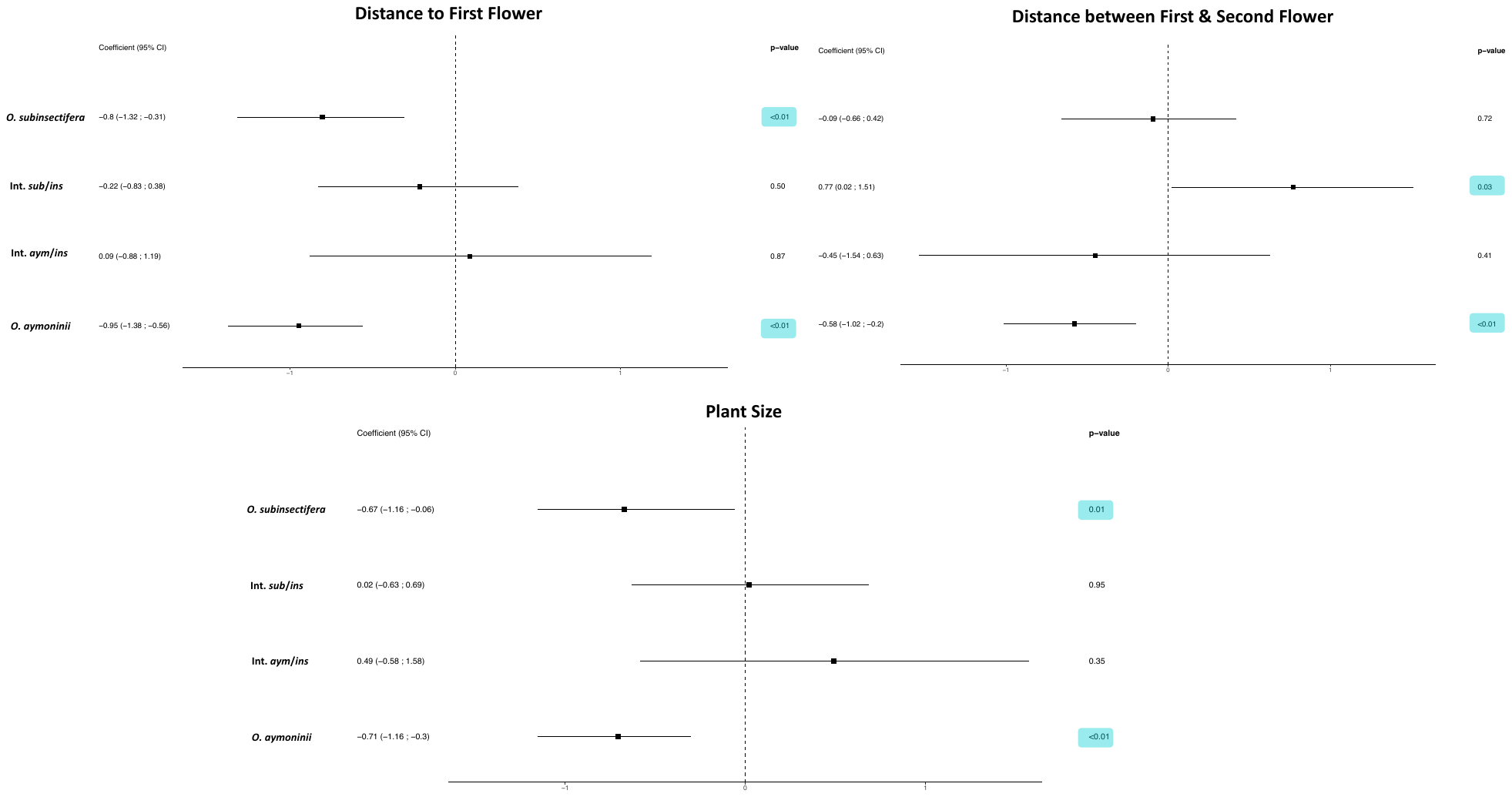

**Figure S2.** Forest plots of the different Floral and Whole-plant traits measured for each species and their intermediates. Significant *p*-values are highlighted in blue.

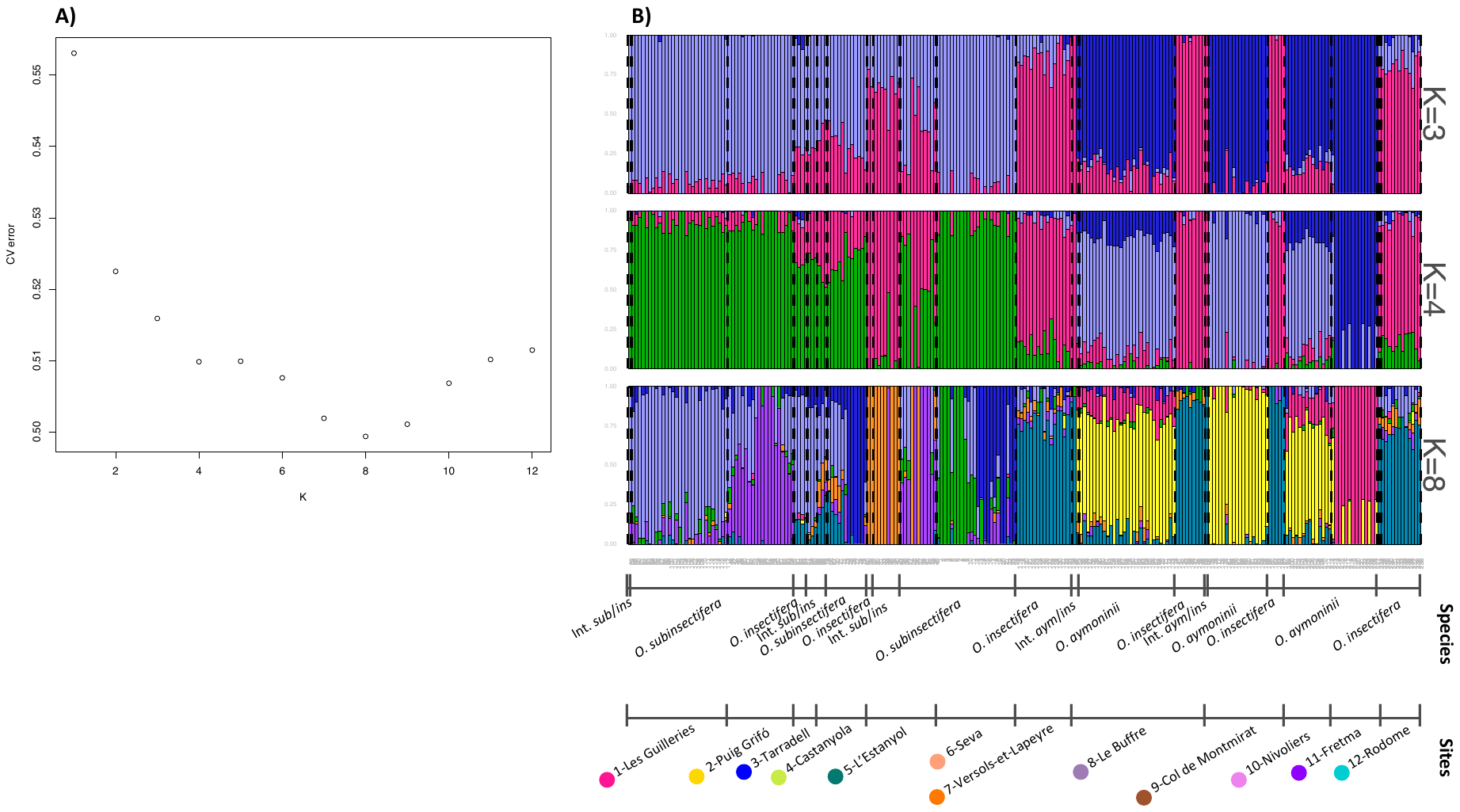

**Figure S3. A** Values of the cross-entropy criterion for a number of clusters ranging from *K*=1 to 12. The optimal number of *K* was found to be 8. **B** Barplot of ancestry coefficients obtained from ADMIXTURE for 239 individuals for *K*=3, *K*=4 and *K*=8, based on 11,089 SNPs.

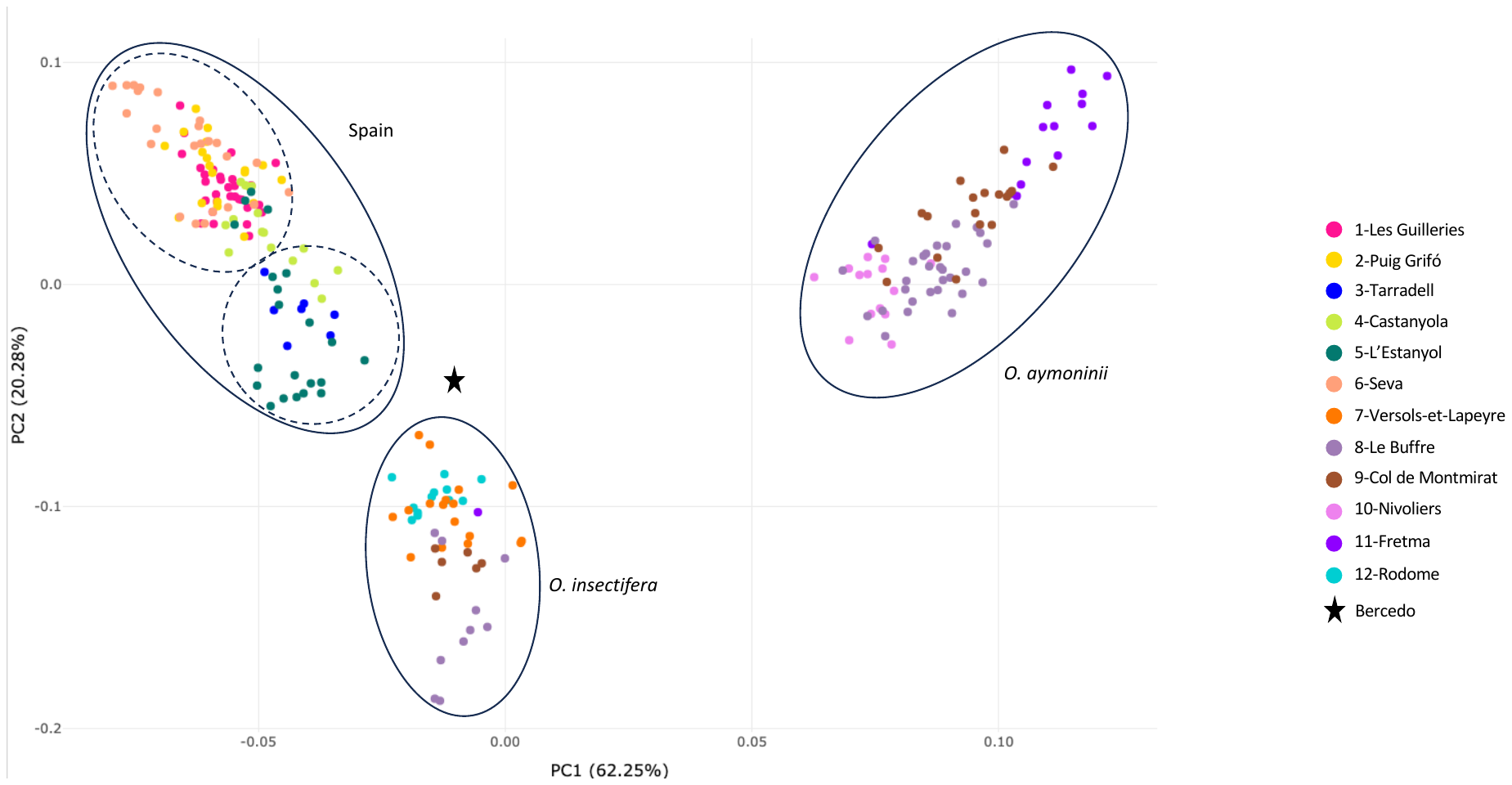

**Figure S4.** Principal Component Analysis (PCA) displaying the two first axes (PC1 and PC2) representing 62.25% and 20.28% of the total genetic variance. PCA was computed on 240 individuals from the 12 sites included in this study and the site Bercedo genotyped using the same protocol (Gibert *et al*., 2023) at 11,534 SNPs and colors depict sampling localities.

**Figure S5.** Phylogenetic tree generated using Astral-III (Zhang *et al*., 2018) based on the concatenation of 4727 locus trees generated with IQ-TREE (Minh *et al*., 2020) with the best model GTR + I + G. The sites (3-Tarradell, 4-Castanyola and 5-L’Estanyol) with intermediate individuals were removed from the analysis. The three species *O. aymoninii*, *O. subinsectifera* and *O. insectifera* were found to be monophyletic with a bootstrap of 1, with the exception of two individuals 22-Div-220 and 23-Div-230, which were not phenotypically different but might be cryptically introgressed.
